# Fast Bayesian Inference of Copy Number Variants using Hidden Markov Models with Wavelet Compression

**DOI:** 10.1101/023705

**Authors:** John Wiedenhoeft, Eric Brugel, Alexander Schliep

## Abstract

By combining Haar wavelets with Bayesian Hidden Markov Models, we improve detection of genomic copy number variants (CNV) in array CGH experiments compared to the state-of-the-art, including standard Gibbs sampling. At the same time, we achieve drastically reduced running times, as the method concentrates computational effort on chromosomal segments which are difficult to call, by dynamically and adaptively recomputing consecutive blocks of observations likely to share a copy number. This makes routine diagnostic use and re-analysis of legacy data collections feasible; to this end, we also propose an effective automatic prior. An open source software implementation of our method is available at http://bioinformatics.rutgers.edu/Software/HaMMLET/. The web supplement is at http://bioinformatics.rutgers.edu/Supplements/HaMMLET/.

**Author Summary:** Identifying large-scale genome deletions and duplications, or copy number variants (CNV), accurately in populations or individual patients is a crucial step in indicating disease factors or diagnosing an individual patient's disease type. Hidden Markov Models (HMM) are a type of statistical model widely used for CNV detection, as well as other biological applications such as the analysis of gene expression time course data or the analysis of discrete-valued DNA and protein sequences.

As with many statistical models, there are two fundamentally different inference approaches. In the frequentist framework, a single estimate of the model parameters would be used as a basis for subsequent inference, making the identification of CNV dependent on the quality of that estimate. This is an acute problem for HMM as methods for finding globally optimal parameters are not known. Alternatively, one can use a Bayesian approach and integrate over all possible parameter choices. While the latter is known to lead to significantly better results, the much—up to hundreds of times—larger computational effort prevents wide adaptation so far.

Our proposed method addresses this by combining Haar wavelets and HMM. We greatly accelerate fully Bayesian HMMs, while simultaneously increasing convergence
and thus the accuracy of the Gibbs sampler used for Bayesian computations, leading to substantial improvements over the state-of-the-art.

## Introduction

The human genome shows remarkable plasticity, leading to significant copy number variations (CNV) within the human population [1]. They contribute to differences in phenotype [2–4], ranging from benign variation over disease susceptibility to inherited and somatic diseases [5], including neuropsychiatric disorders [6–8] and cancer [9,10]. Separating common from rare variants is important in the study of genetic diseases [5, 11, 12], and while the experimental platforms have matured, interpretation and assessment of pathogenic significance remains a challenge [13].

Computationally, CNV detection is a segmentation problem, in which consecutive stretches of the genome are to be labeled by their copy number; following the conventions typically employed in CNV method papers, e.g. [14–17], we use this term rather abstractly to denote segments of equal mean value, not actual ploidy, though for homogeneous samples the latter can be easily assigned. Along with a variety of other methods [14–16,18–41], Hidden Markov Models (HMM) [42] play a central role [17,43–52], as they directly model the separate layers of observed measurements, such as log-ratios in array comparative genomic hybridization (aCGH), and their corresponding latent copy number (CN) states, as well as the underlying linear structure of segments.

As statistical models, they depend on a large number of parameters, which have to be either provided *a priori* by the user or inferred from the data. Classic frequentist maximum likelihood (ML) techniques like Baum-Welch [53,54] are not guaranteed to be globally optimal, i. e. they can converge to the wrong parameter values, which can limit the accuracy of the segmentation. Furthermore, the Viterbi algorithm [55] only yields a single maximum a posteriori (MAP) segmentation given a parameter estimate [56]. Failure to consider the full set of possible parameters precludes alternative interpretations of the data, and makes it very difficult to derive p-values or confidence intervals. Furthermore, these frequentist techniques have come under increased scrutiny in the scientific community.

Bayesian inference techniques for HMMs, in particular Forward-Backward Gibbs sampling [57,58], provide an alternative for CNV detection as well [59–61]. Most importantly, they yield a complete probability distribution of copy numbers for each observation. As they are sampling-based, they are computationally expensive, which is problematic especially for high-resolution data. Though they are guaranteed to converge to the correct values under very mild assumptions, they tend to do so slowly, which can lead to oversegmentation and mislabeling if the sampler is stopped prematurely.

Another issue in practice is the requirement to specify hyperparameters for the prior distributions. Despite the theoretical advantage of making the inductive bias more explicit, this can be a major source of annoyance for the user. It is also hard to justify any choice of hyperparameters when insufficient domain knowledge is available.

Recent work of our group [62] has focused on accelerating Forward-Backward Gibbs sampling through the introduction of compressed HMMs and approximate sampling. For the first time, Bayesian inference could be performed at running times on par with classic maximum likelihood approaches. It was based on a greedy spatial clustering heuristic, which yielded a static compression of the data into blocks, and block-wise sampling. Despite its success, several important issues remain to be addressed. The blocks are fixed throughout the sampling and impose a structure that turns out to be too rigid if variances differ between CN states. The clustering heuristic relies on empirically derived parameters not supported by a theoretical analysis, which can lead to suboptimal clustering or overfitting. Also, the method cannot easily be generalized for multivariate data. Lastly, the implementation was primarily aimed at comparative analysis between the frequentist and Bayesian approach, as opposed to overall speed.

To address these issues and make Bayesian CNV inference feasible even on a laptop, we propose the combination of HMMs with another popular signal processing technology: Haar wavelets have previously been used in CNV detection [63], mostly as a preprocessing tool for statistical downstream applications [28–32] or simply as a visual aid in GUI applications [21,64]. A combination of smoothing and segmentation has been suggested as likely to improve results [65], and here we show that this is indeed the case. Wavelets provide a theoretical foundation for a better, dynamic compression scheme for faster convergence and accelerated Bayesian sampling. We improve simultaneously upon the typically conflicting goals of accuracy and speed, because the wavelets allow summary treatment of “easy” CN calls in segments and focus computational effort on the “difficult” CN calls, dynamically and adaptively. This is in contrast to other computationally efficient tools, which often simplify the statistical model or use heuristics. The required data structure can be efficiently computed, incurs minimal overhead, and has a straightforward generalization for multivariate data. We further show how the wavelet transform yields a natural way to set hyperparameters automatically, with little input from the user.

We implemented our method in a highly optimized end-user software, called HaMMLET. Aside from achieving an acceleration of up to two orders of magnitude, it exhibits significantly improved convergence behavior, has excellent precision and recall, and provides Bayesian inference within seconds even for large data sets. The accuracy and speed of HaMMLET also makes it an excellent choice for routine diagnostic use and large-scale re-analysis of legacy data. Notice that while we focus on aCGH in this paper as the most straightforward biological example of univariate Gaussian data, the method we present is a general approach to Bayesian HMM inference as long as the emission distributions come from the exponential family, implying that conjugate priors exist and the dimension of its sufficient statistics remain bounded with increasing sample size. It can thus be readily generalized and adapted to read-depth data, SNP arrays, and multi-sample applications.

## Results

### Simulated aCGH data

A previous survey [65] of eleven CNV calling methods for aCGH has established that segmentation-focused methods such as DNAcopy [14, 36], an implementation of circular binary segmentation (CBS), as well as CGHseg [37] perform consistently well. DNAcopy performs a number of t-tests to detect break-point candidates. The result is typically oversegmented and requires a merging step in post-processing, especially to reduce the number of segment means. To this end MergeLevels was introduced by [66]. They compare the combination DNAcopy+MergeLevels to their own HMM implementation [17] as well as GLAD (“Gain and Loss Analysis of DNA”) [27], showing its superior performance over both methods. This established DNAcopy+MergeLevels as the *de facto* standard in CNV detection, despite the comparatively long running time.

The paper also includes aCGH simulations deemed to be reasonably realistic by the community. DNACopy was used to segment 145 unpublished samples of breast cancer data, and subsequently labeled as copy numbers 0 to 5 by sorting them into bins with boundaries (–∞, –0.4, –0.2, 0.2,0.4,0.6, ∞), based on the sample mean in each segment (the last bin appears to not be used). Empirical length distributions were derived, from which the sizes of CN aberrations are drawn. The data itself is modeled to include Gaussian noise, which has been established as sufficient for aCGH data [67]. Means were generated such as to mimic random tumor cell proportions, and random variances were chosen to simulate experimenter bias often observed in real data; this emphasizes the importance of having automatic priors available when using Bayesian methods, as the means and variances might be unknown *a priori*. The data comprises three sets of simulations: “breakpoint detection and merging” (BD&M), “spatial resolution study” (SRS), and “testing” (T) (see their paper for details). We used the MergeLevels implementation as provided on their website. It should be noted that the superiority of DNAcopy+MergeLevels was established using a simulation based upon segmentation results of DNAcopy itself.

We used the Bioconductor package DNAcopy (version 1.24.0), and followed the procedure suggested therein, including outlier smoothing. This version uses the linear-time variety of CBS [15]; note that other authors such as [35] compare against a quadratic-time version of CBS [14], which is significantly slower. For HaMMLET, we use a 5-state model with automatic hyperparameters ℙ(σ^2^ ≤ 0.01) = 0.9 (see section Automatic priors), and all Dirichlet hyperparameters set to 1.

Following [62], we report F-measures (*F*_1_ scores) for binary classification into normal and aberrant segments (Fig. 2), using the usual definition of 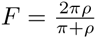 being the harmonic mean of precision 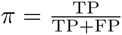 and recall 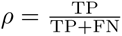, where TP, FP, TN and FN denote true/false positives/negatives, respectively. On datasets T and BD&M, both methods have similar medians, but HaMMLET has a much better interquartile range (IQR) and range, about half of CBS’s. On the spatial resolution data set (SRS), HaMMLET performs much better on very small aberrations. This might seem somewhat surprising, as short segments could easily get lost under compression. However, Lai *et al*. [65] have noted that smoothing-based methods such as quantile smoothing (quantreg) [23], lowess [24], and wavelet smoothing [29] perform particularly well in the presence of high noise and small CN aberrations, suggesting that “an optimal combination of the smoothing step and the segmentation step may result in improved performance”. Our wavelet-based compression inherits those properties. For CNVs of sizes between 5 and 10, CBS and HaMMLET have similar ranges, with CBS being skewed towards better values; CBS has a slightly higher median for 10–20, with IQR and range being about the same. However, while HaMMLET’s F-measure consistently approaches 1 for larger aberrations, CBS does not appear to significantly improve after size 10. The plots for all individual samples can be found in Web Supplement S1–S3, which can be viewed online at http://schlieplab.org/Supplements/HaMMLET/, or downloaded from https://zenodo.org/record/46263(DOI: 10.5281/zenodo.46263).

**Figure 1.**
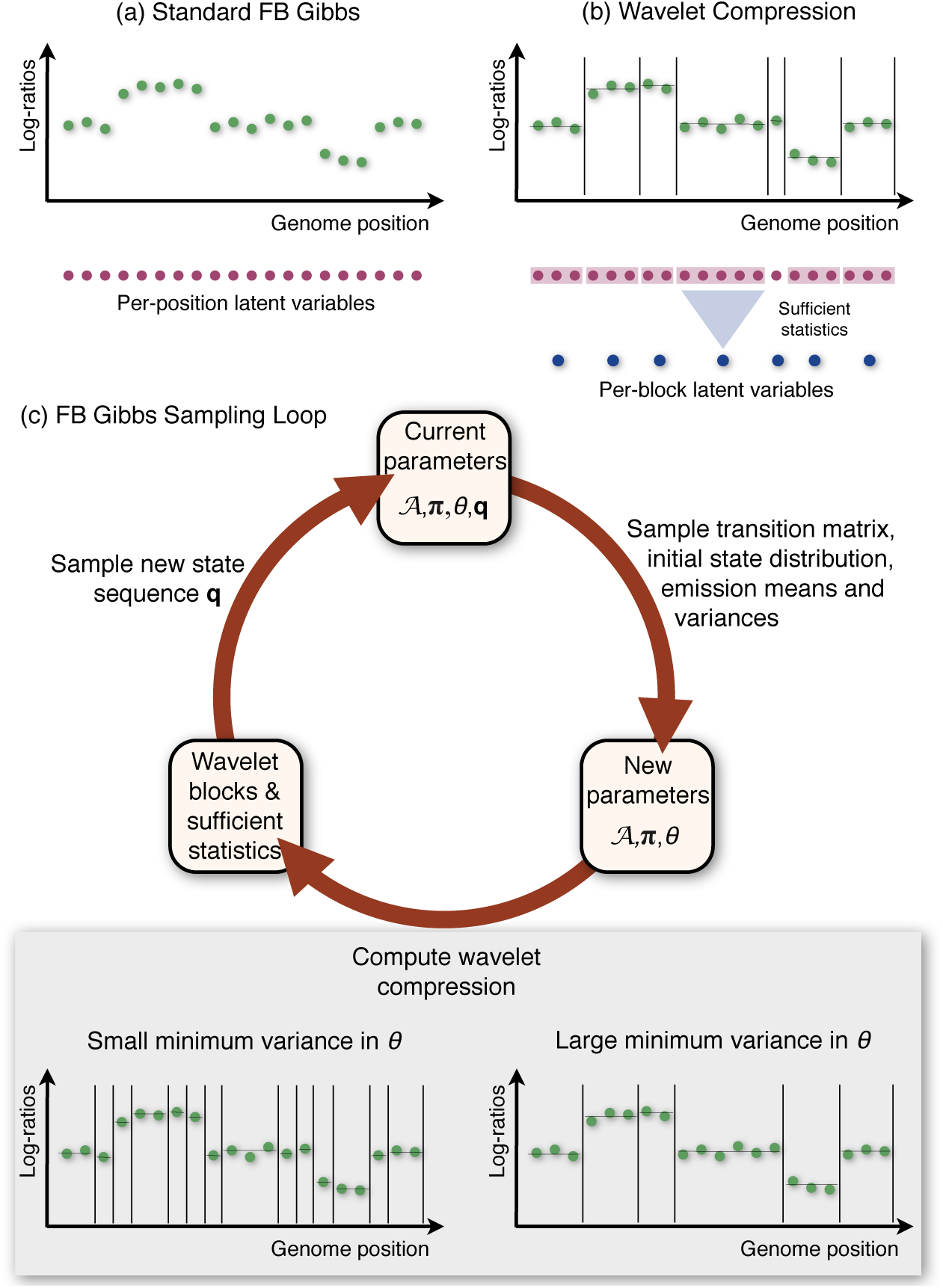
Overview of HaMMLET. Istead of individual computations per observation (panel a), Forward-Backward Gibbs Sampling is performed on a compressed version of the data, using sufficient statistics for block-wise computations (panel b) to accelerate inference in Bayesian Hidden Markov Models. During the sampling (panel c) parameters and copy number sequences are sampled iteratively. During each iteration, the sampled emission variances determine which coefficients of the data’s Haar wavelet transform are dynamically set to zero. This controls potential break points at finer or coarser resolution or, equivalently, defines blocks of variable number and size (panel c, bottom). Our approach thus yields a dynamic, adaptive compression scheme which greatly improves speed of convergence, accuracy and running times.

**Figure 2.**
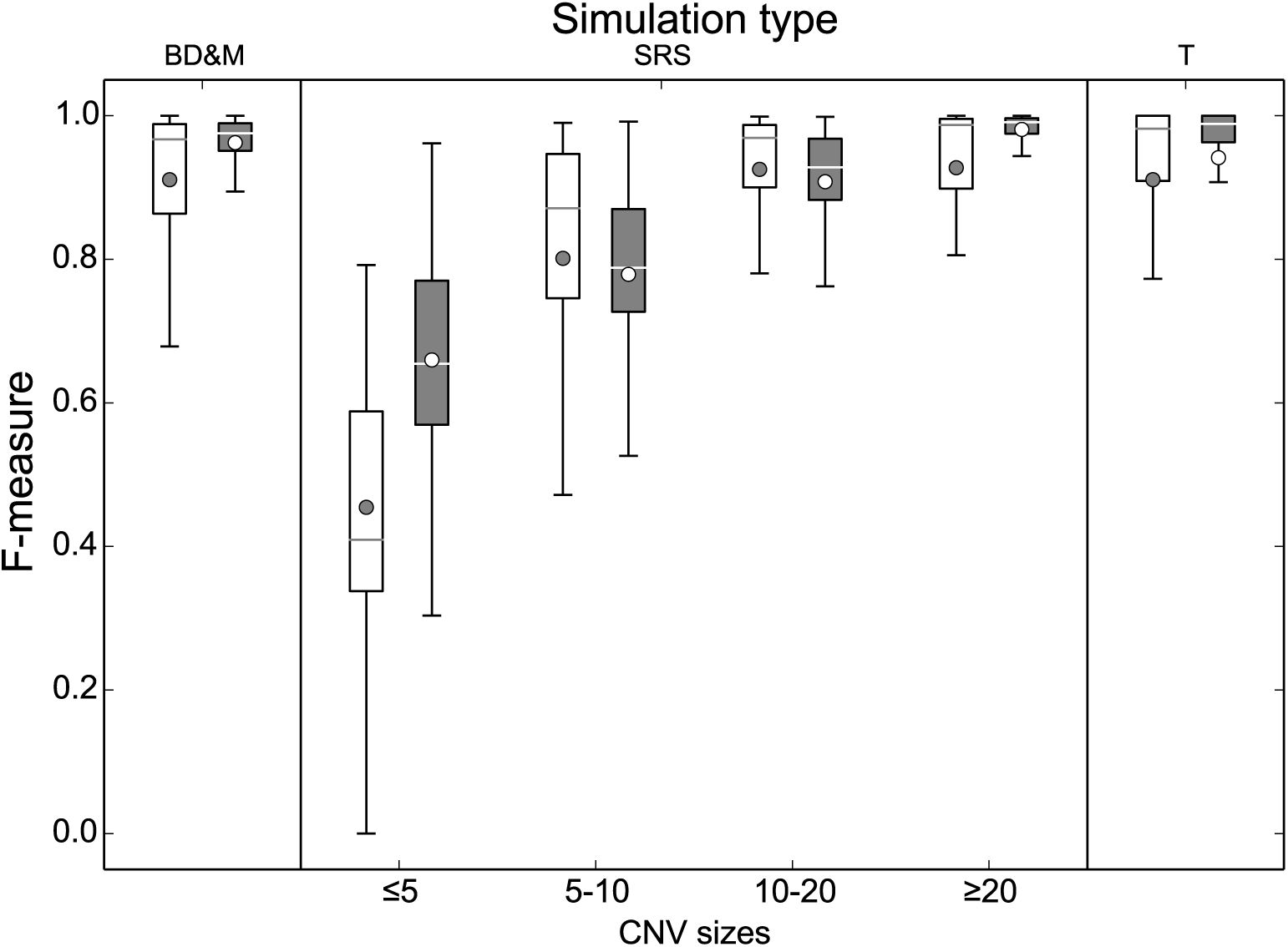
F-measures of CBS (light) and HaMMLET (dark) for calling aberrant copy numbers on simulated aCGH data [66]. Boxes represent the interquartile range (IQR = Q3-Q1), with a horizontal line showing the median (Q2), whiskers representing the range (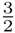IQR beyond Q1 and Q3), and the bullet representing the mean. HaMMLET has the same or better F-measures in most cases, and on the SRS simulation converges to 1 for larger segments, whereas CBS plateaus for aberrations greater than 10.

### High-density CGH array

In this section, we demonstrate HaMMLET’s performance on biological data. Due to the lack of a gold standard for high-resolution platforms, we assess the CNV calls qualitatively. We use raw aCGH data (GEO:GSE23949) [68] of genomic DNA from breast cancer cell line BT-474 (invasive ductal carcinoma, GEO:GSM590105), on an Agilent-021529 Human CGH Whole Genome Microarray 1x1M platform (GEO:GPL8736). We excluded gonosomes, mitochondrial and random chromosomes from the data, leaving 966,432 probes in total.

HaMMLET allows for using automatic emission priors (see section Automatic priors) by specifying a noise variance, and a probability to sample a variance not exceeding this value. We compare HaMMLET’s performance against CBS, using a 20-state model with automatic priors, ℙ(σ^2^ ≤ 0.1) = 0.8, 10 prior self-transitions and 1 for all other hyperparameters. CBS took over 2h9m to process the entire array, whereas HaMMLET took 27.1s for 100 iterations, a speedup of 288. The compression ratio (see section Effects of wavelet compression on speed and convergence) was 220.3. CBS yielded a massive over-segmentation into 1,548 different copy number levels; cf. Supplement S4 at https://zenodo.org/record/46263. As the data is derived from a relatively homogeneous cell line as opposed to a biopsy, we do not expect the presence of subclonal populations to be a contributing factor [69,70]. Instead, measurements on aCGH are known to be spatially correlated, resulting in a wave pattern which has to be removed in a preprocessing step; notice that the internal compression mechanism of HaMMLET is derived from a spatially adaptive regression method, so smoothing is inherent to our method. CBS performs such a smoothing, yet an unrealistically large number of different levels remains, likely due to residuals of said wave pattern. Furthermore, repeated runs of CBS yielded different numbers of levels, suggesting that indeed the merging was incomplete. This can cause considerable problems downstream, as many methods operate on labeled data. A common approach is to consider a small number of classes, typically 3 to 4, and associate them semantically with CN labels like *loss, neutral, gain*, and *amplification*, e.g. [27,59,61,67,71–75]. In inference models that contain latent categorical state variables, like HMM, such an association is readily achieved by sorting classes according to their means. In contrast, methods like CBS typically yield a large, often unbounded number of classes, and reducing it is the declared purpose of merging algorithms, see [66]. Consider, for instance, CGHregions [74], which uses a 3-label matrix to define regions of shared CNV events across multiple samples by requiring a maximum *L*_1_ distance of label signatures between all probes in that region. If the domain of class labels was unrestricted and potentially different in size for each sample, such a measure would not be meaningful, since the *i*-th out of *n* classes cannot be readily identified with the *i*-th out of *m* classes for *n* ≠ *m*, hence no two classes can be said to represent the same CN label. Similar arguments hold true for clustering based on Hamming distance [72] or ordinal similarity measures [71]. Furthermore, even CGHregions’ optimized computation of medoids takes several minutes to compute. As the time depends multiplicatively on the number of labels, increasing it by three orders of magnitude would increase downstream running times to many hours.

For a more comprehensive analysis, we restricted our evaluation to chromosome 20 (21,687 probes), which we assessed to be the most complicated to infer, as it appears to have the highest number of different CN states and breakpoints. CBS yields a 19-state result after 15.78s (Fig. 3, top). We have then used a 19-state model with automated priors (ℙ(σ^2^ ≤ 0.04) = 0.9), 10 prior self-transitions, all other Dirichlet parameters set to 1) to reproduce this result. Using noise control (see Methods), our method took 1.61s for 600 iterations. The solution we obtained is consistent with CBS (Fig. 3, middle and bottom). However, only 11 states were part of the final solution, i.e. 8 states yielded no significant likelihood above that of other states. We observe superfluous states being ignored in our simulations as well. In light of the results on the entire array, we suggest that the segmentation by DNAcopy has not sufficiently been merged by MergeLevels. Most strikingly, HaMMLET does not show any marginal support for a segment called by CBS around probe number 4,500. We have confirmed that this is not due to data compression, as the segment is broken up into multiple blocks in each iteration (cf. Supplement S5 at https://zenodo.org/record/46263). On the other hand, two much smaller segments called by CBS in the 17,000-20,000 range do have marginal support of about 40% in HaMMLET, suggesting that the lack of support for the larger segment is correct. It should be noted that inference differs between the entire array and chromosome 20 in both methods, since long-range effects have higher impact in larger data.

**Figure 3.**
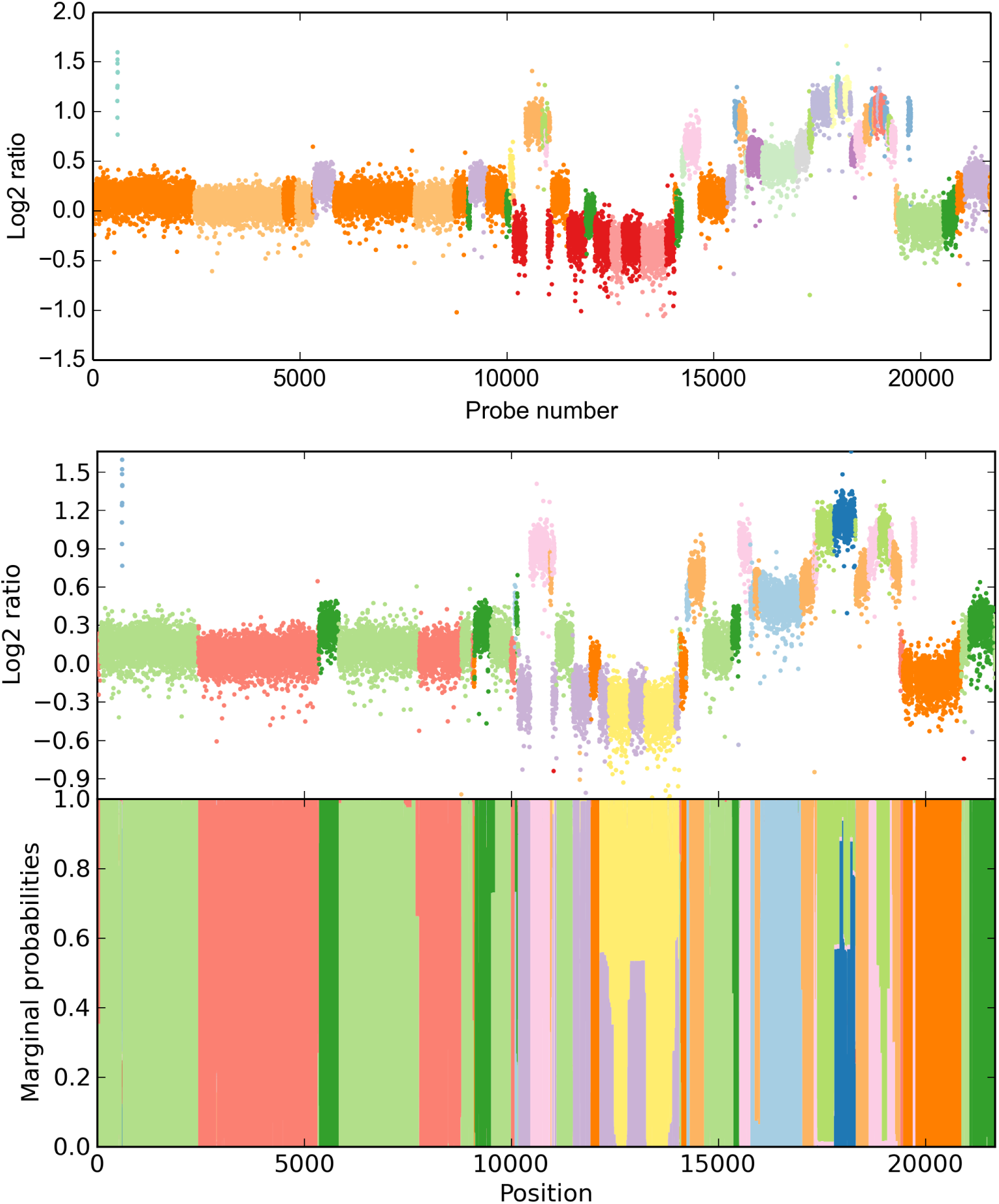
Copy number inference for chromosome 20 in invasive ductal carcinoma (21,687 probes) CBS creates a 19-state solution (top), however, a compressed 19-state HMM only supports an 11-state solution (bottom), suggesting insufficient level merging in CBS. Also notice the additional diagnostic gain provided by the marginal probabilities, allowing to assess uncertainty in annotation, e.g. around position 13,000.

We also demonstrate another feature of HaMMLET called *noise control*. While Gaussian emissions have been deemed a sufficiently accurate noise model for aCGH [67], microarray data is prone to outliers, for example due to damages on the chip. While it is possible to model outliers directly [60], the characteristics of the wavelet transform allow us to largely suppress them during the construction of our data structure (see Methods). Notice that due to noise control most outliers are correctly labeled according to the segment they occur in, while the short gain segment close to the beginning is called correctly.

### Effects of wavelet compression on speed and convergence

The speedup gained by compression depends on how well the data can be compressed. Poor compression is expected when the means are not well separated, or short segments have small variance, which necessitates the creation of smaller blocks for the rest of the data to expose potential low-variance segments to the sampler. On the other hand, data must not be over-compressed to avoid merging small aberrations with normal segments, which would decrease the F-measure. Due to the dynamic changes to the block structure, we measure the level of compression as the average compression ratio, defined as the product of the number of data points *T* and the number of iterations *N*, divided by the total number of blocks in all iterations. As usual a compression ratio of 1 indicates no compression.

To evaluate the impact of dynamic wavelet compression on speed and convergence properties of an HMM, we created 129,600 different data sets with *T* = 32,768 many probes. In each data set, we randomly distributed 1 to 6 gains of a total length of {100, 250, 500, 750, 1000 } uniformly among the data, and do the same for losses. Mean combinations (*μ*_loss_,*μ*_neutral_,*μ*_gain_) were chosen from 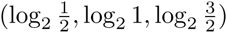, (–1,0,1), (–2,0,2), and (–10,0,10), and variances 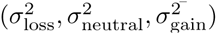 from (0.05,0.05,0.05), (0.5,0.1,0.9), (0.3,0.2,0.1), (0.2,0.1,0.3), (0.1,0.3,0.2), and (0.1,0.1,0.1). These values have been selected to yield a wide range of easy and hard cases, both well separated, low-variance data with large aberrant segments as well as cases in which small aberrations overlap significantly with the tail samples of high-variance neutral segments. Consequently, compression ratios range from ~1 to ~2,100. We use automatic priors ℙ(σ^2^ ≤ 0.2) = 0.9, self-transition priors *α_ii_* ∈ {10,100,1000 }, non-self transition priors *α_ij_* = 1, and initial state priors ***α*** ∈ {**1,10** }. Using all possible combinations of the above yields 129,600 different simulated data sets, a total of 4.2 billion values.

We achieve speedups per iteration of up to 350 compared to an uncompressed HMM (Fig. 4). In contrast, [62] have reported ratios of 10–60, with one instance of 90. Notice that the speedup is not linear in the compression ratio. While sampling itself is expected to yield linear speedup, the marginal counts still have to be tallied individually for each position, and dynamic block creation causes some overhead. The quantization artifacts observed for larger speedup are likely due to the limited resolution of the Linux time command (10 milliseconds). Compressed HaMMLET took about 11.2 CPU hours for all 129,600 simulations, whereas the uncompressed version took over 3 weeks and 5 days. All running times reported are CPU time measured on a single core of a AMD Opteron 6174 Processor, clocked at 2.2 GHz.

**Figure 4.**
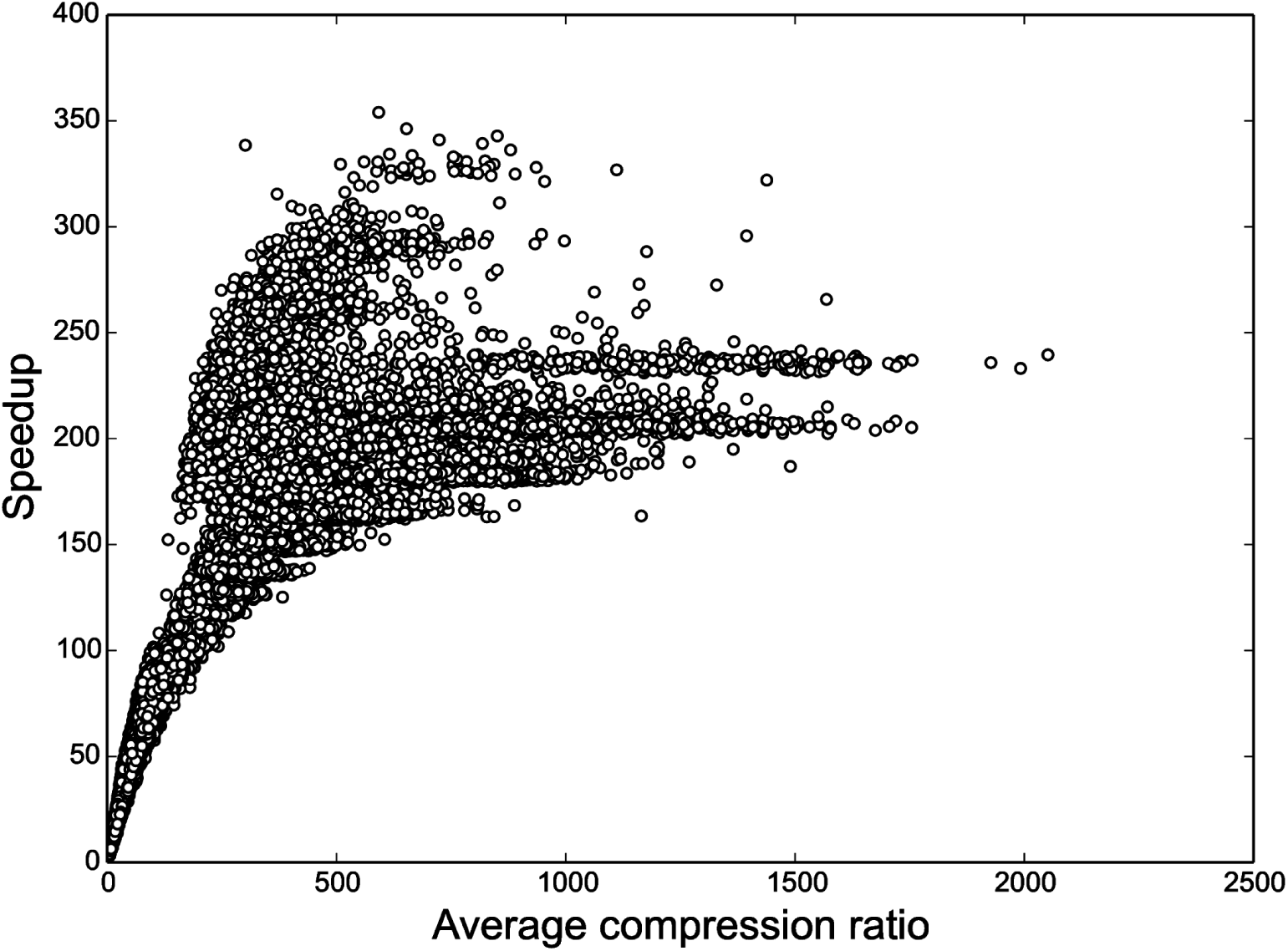
HaMMLET’s speedup as a function of the average compression during sampling. As expected, higher compression leads to greater speedup. The non-linear characteristic is due to the fact that some overhead is incurred by the dynamic compression, as well as parts of the implementation that do not depend on the compression, such as tallying marginal counts.

We evaluate the convergence of the F-measure of compressed and uncompressed inference for each simulation. Since we are dealing with multi-class classification, we use the micro - and macro-averaged F-measures (*F*_mi_, *F*_ma_) proposed by [76],

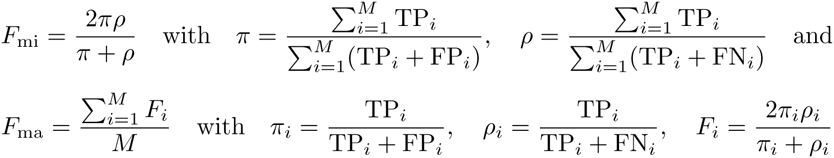

Here, *TP_i_* denotes a true positive call for the *i*-th out of *M* states, *π* and *ρ* denote precision and recall. These F-measures tend to be dominated by the classifier’s performance on common and rare categories, respectively. Since all state labels are sampled from the same prior and hence their relative order is random, we used the label permutation which yielded the highest sum of micro - and macro-averaged F-measures. The simulation results are included in Supplement S6 at https://zenodo.org/record/46263.

In Fig. 5, we show that the compressed version of the Gibbs sampler converges almost instantly, whereas the uncompressed version converges much slower, with about 5% of the cases failing to yield an F-measure > 0.6 within 1,000 iterations. Wavelet compression is likely to yield reasonably large blocks for the majority class early on, which leads to a strong posterior estimate of its parameters and self-transition probabilities. As expected, *F*_ma_ are generally worse, since any misclassification in a rare class has a larger impact. Especially in the uncompressed version, we observe that *F*_ma_ tends to plateau until *F*_mi_ approaches 1.0. Since any misclassification in the majority (neutral) class adds false positives to the minority classes, this effect is expected. It implies that correct labeling of the majority class is a necessary condition for correct labeling of minority classes, in other words, correct identification of the rare, interesting segments requires the sampler to properly converge, which is much harder to achieve without compression. It should be noted that running compressed HaMMLET for 1,000 iterations is unnecessary on the simulated data, as in all cases it converges between 25 and 50 iterations. Thus, for all practical purposes, further speedup by a factor of 40–80 can be achieved by reducing the number of iterations, which yields convergence up to 3 orders of magnitude faster than standard FBG.

**Figure 5.**
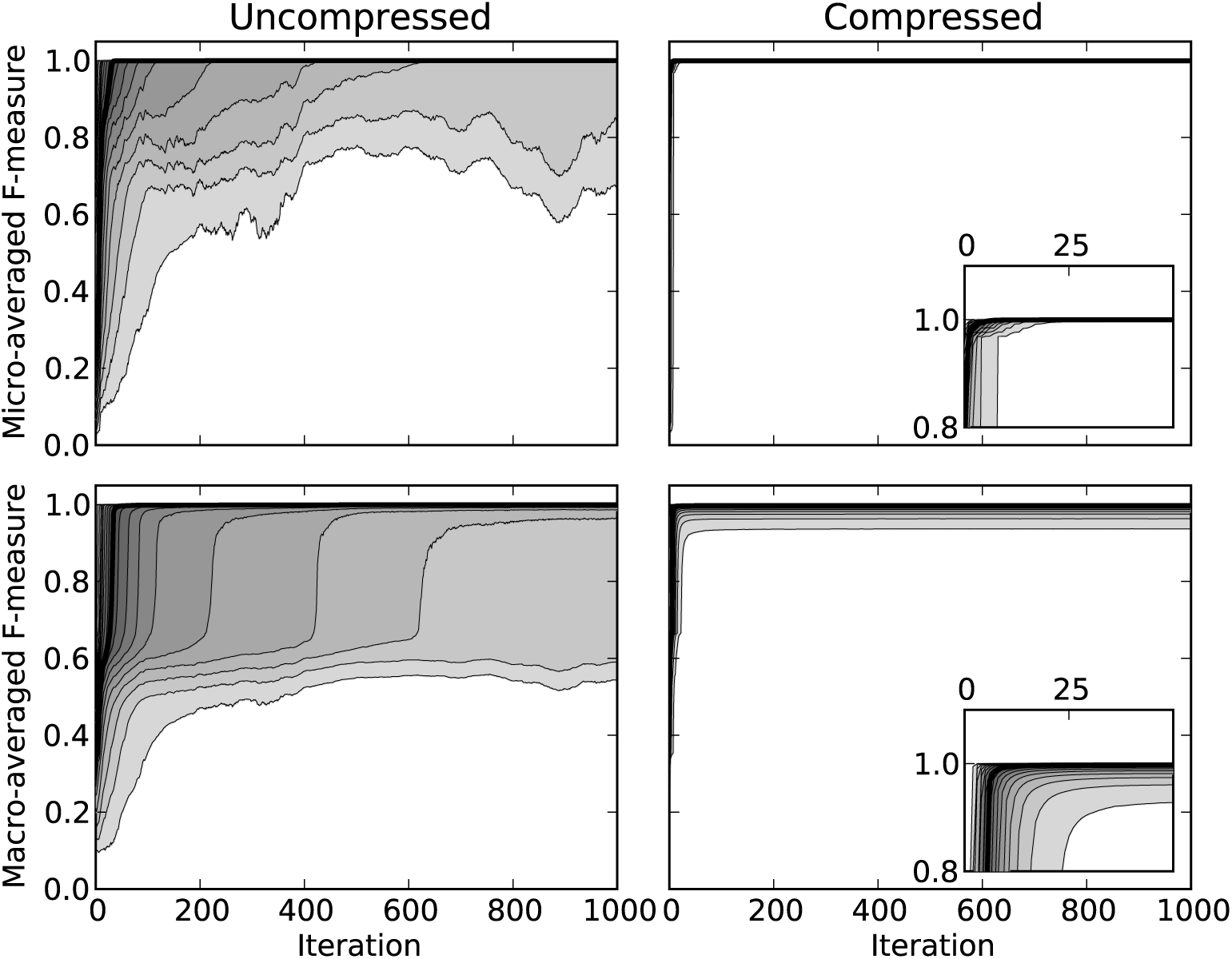
F-measures for simulation results. The median value (black) and quantile ranges (in 5% steps) of the micro - (top) and macro-averaged (bottom) F-measures (*F*_mi_, *F*_ma_) for uncompressed (left) and compressed (right) FBG inference, on the same 129,600 simulated data sets, using automatic priors. The x-axis represents the number of iterations alone, and does not reflect the additional speedup obtained through compression. Notice that the compressed HMM converges no later than 50 iterations (inset figures, right).

### Coriell, ATCC and breast carcinoma

The data provided by [77] includes 15 aCGH samples for the Coriell cell line. At about 2,000 probes, the data is small compared to modern high-density arrays. Nevertheless, these data sets have become a common standard to evaluate CNV calling methods, as they contain few and simple aberrations. The data also contains 6 ATCC cell lines as well as 4 breast carcinoma, all of which are substantially more complicated, and typically not used in software evaluations. In Fig. 6, we demonstrate our ability to infer the correct segments on the most complex example, a T47D breast ductal carcinoma sample of a 54 year old female. We used 6-state automatic priors with ℙ(σ^2^ ≤ 0.1) = 0.85, and all Dirichlet hyperparameters set to 1. On a standard laptop, HaMMLET took 0.09 seconds for 1,000 iterations; running times for the other samples were similar. Our results for all 25 data sets have been included in Supplement S7 at https://zenodo.org/record/46263.

**Figure 6.**
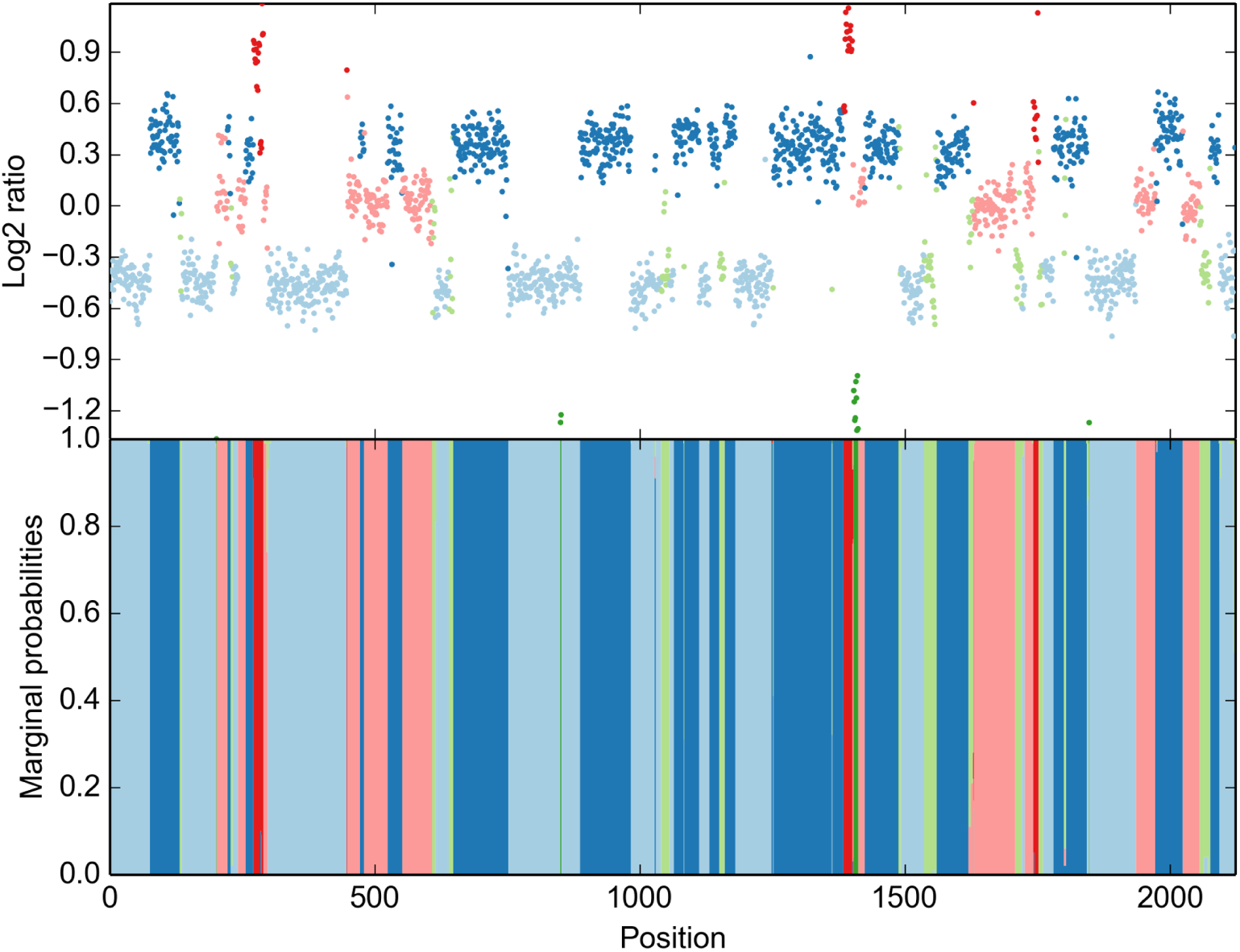
HaMMLET’s inference of copy-number segments on T47D breast ductal carcinoma. Notice that the data is much more complex than the simple structure of a diploid majority class with some small aberrations typically observed for Coriell data.

## Discussion

In the analysis of biological data, there are usually conflicting objectives at play which need to be balanced: the required accuracy of the analysis, ease of use—using the software, setting software and method parameters—and often the speed of a method.

Bayesian methods have obtained a reputation of requiring enormous computational effort and being difficult to use, for the expert knowledge required for choosing prior distributions. It has also been recognized [60,62,78] that they are very powerful and accurate, leading to improved, high-quality results and providing, in the form of posterior distributions, an accurate measure of uncertainty in results. Nevertheless, it is not surprising that a hundred times larger effort in computation alone prevented wide-spread use.

Inferring Copy Number Variants (CNV) is a quite special problem, as experts can identify CN changes visually, at least on very good data and for large, drastic CN changes (e. g., long segments lost on both chromosomal copies). With lesser quality data, smaller CN differences and in the analysis of cohorts the need for objective, highly accurate, and automated methods is evident.

The core idea for our method expands on our prior work [62] and affirms a conjecture by Lai *et al*. [65] that a combination of smoothing and segmentation will likely improve results. One ingredient of our method are Haar wavelets, which have previously been used for pre-processing and visualization [21,64]. In a sense, they quantify and identify regions of high variation, and allow to summarize the data at various levels of resolution, somewhat similar to how an expert would perform a visual analysis. We combine, for the first time, wavelets with a full Bayesian HMM by dynamically and adaptively infering blocks of subsequent observations from our wavelet data structure. The HMM operates on blocks instead of individual observations, which leads to great saving in running times, up to 350-fold compared to the standard FB-Gibbs sampler, and up to 288 times faster than CBS. Much more importantly, operating on the blocks greatly improves convergence of the sampler, leading to higher accuracy for a much smaller number of sampling iterations. Thus, the combination of wavelets and HMM realizes a simultaneous improvement on accuracy and on speed; typically one can have one or the other. An intuitive explanation as to why this works is that the blocks derived from the wavelet structure allow efficient, summary treatment of those “easy” to call segments given the current sample of HMM parameters and identifies “hard” to call CN segment which need the full computational effort from FB-Gibbs. Note that it is absolutely crucial that the block structure depends on the parameters sampled for the HMM and will change drastically over the run time. This is in contrast to our prior work [62], which used static blocks and saw no improvements to accuracy and convergence speed. The data structures and linear-time algorithms we introduce here provide the efficient means for recomputing these blocks at every cycle of the sampling, cf. Fig. 1. Compared to our prior work, we observe a speed-up of up to 3,000 due to the greatly improved convergence, *O*(*T*) vs. *O*(*T* log *T*) clustering, improved numerics and, lastly, a C++ instead of a Python implementation.

We performed an extensive comparison with the state-of-the-art as identified by several review and benchmark publications and with the standard FB-Gibbs sampler on a wide range of biological data sets and 129,600 simulated data sets, which were produced by a simulation process not based on HMM to make it a harder problem for our method. All comparisons demonstrated favorable results for our method when measuring accuracy at a very noticeable acceleration compared to the state-of-the-art. It must be stressed that these results were obtained with a statistically sophisticated model for CNV calls and without cutting algorithmic corners, but rather an effective allocation of computational effort.

All our computations are performed using our automatic prior, which derives estimates for the hyperparameters of the priors for means and variances directly from the wavelet tree structure and the resulting blocks. The block structure also imposes a prior on self-transition probabilities. The user only has to provide an estimate of the noise variance, but as the automatic prior is designed to be weak, the prior and thus the method is robust against incorrect estimates. We have demonstrated this by using different hyperparameters for the associated Dirichlet priors in our simulations, which HaMMLET is able to infer correctly regardless of the transition priors. At the same time the automatic prior can be used to tune certain aspects of the HMM if stronger prior knowledge is available. We would expect further improvements from non-automatic, expert-selected priors, but refrained from using them for the evaluation, as they might be perceived as unfair to other methods.

In summary, our method is as easy to use as other, statistically less sophisticated tools, more accurate and much more computationally efficient. We believe this makes it an excellent choice both for routine use in clinical settings and principled re-analysis of large cohorts, where the added accuracy and the much improved information about uncertainty in copy number calls from posterior marginal distributions will likely yield improved insights into CNV as a source of genetic variation and its relationship to disease.

## Methods

We will briefly review Forward-Backward Gibbs sampling (FBG) for Bayesian Hidden Markov Models, and its acceleration through compression of the data into blocks. By first considering the case of equal emission variances among all states, we show that optimal compression is equivalent to a concept called *selective wavelet reconstruction*, following a classic proof in wavelet theory. We then argue that *wavelet coefficient thresholding*, a variance-dependent minimax estimator, allows for compression even in the case of unequal emission variances. This allows the compression of the data to be adapted to the prior variance level at each sampling iteration. We then derive a simple data structure to dynamically create blocks with little overhead. While wavelet approaches have been used for aCGH data before [29, 33, 34, 63], our method provides the first combination of wavelets and HMMs.

### Bayesian Hidden Markov Models

Let *T* be the length of the observation sequence, which is equal to the number of probes. An HMM can be represented as a statistical model (**q**, ***A**, θ*, **π** | **y**), with transition matrix ***A***, a latent state sequence **q** = (*q*_0_, *q*_1_,…,*q*_*T*–1_), an observed emission sequence **y** = (*y*_0_,*y*_1_, …, *y*_*T*–1_), emission parameters *θ*, and an initial state distribution **π**.

In the usual frequentist approach, the state sequence **q** is inferred by first finding a maximum likelihood estimate of the parameters,

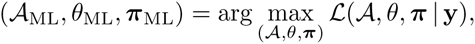

using the Baum-Welsh algorithm [53, 54]. This is only guaranteed to yield local optima, as the likelihood function is not convex. Repeated random reinitializations are used to find “good” local optima, but there are no guarantees for this method. Then, the most likely state sequence given those parameters,

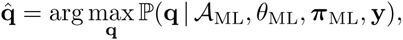

is calculated using the Viterbi algorithm [55]. However, if there are only a few aberrations, that is there is imbalance between classes, the ML parameters tend to overfit the normal state which is likely to yield incorrect segmentation [62]. Furthermore, alternative segmentations given those parameters are also ignored, as are the ones for alternative parameters.

The Bayesian approach is to calculate the distribution of state sequences directly by integrating out the emission and transition variables,

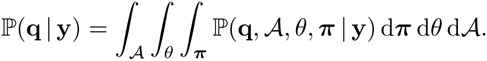

Since this integral is intractable, it has to be approximated using Markov Chain Monte Carlo techniques, i. e. drawing *N* samples,

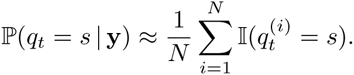

and subsequently approximating marginal state probabilities by their frequency in the sample

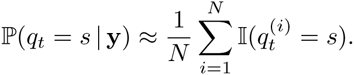

Thus, for each position *t*, we get a complete probability distribution over the possible states. As the marginals of each variable are explicitly defined by conditioning on the other variables, an HMM lends itself to Gibbs sampling, i. e. repeatedly sampling from the marginals (***A*** | **q**, *θ*, **y**, **π**), (*θ* | **q**, ***A***, **y**, **π**), (**π** | ***A**, θ*, **y**, **q**), and (**q** | ***A**, θ*, **y**, **π**), conditioned on the previously sampled values. Using Bayes’ formula and several conditional independence relations, the sampling process can be written as

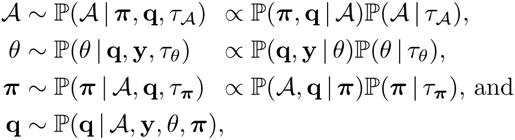

where τ_*x*_ represents hyperparameters to the prior distribution ℙ(*x* | τ_*x*_). Typically, each prior will be conjugate, i. e. it will be the same class of distributions as the posterior, which then only depends on updated parameters τ^⋆^, e.g. 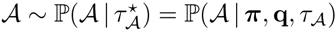. Thus τ_**π**_ and τ_***A***(*k*,:)_, the hyperparameters of **π** and the *k*-th row of ***A***, will be the *α_i_* of a Dirichlet distribution, and τ_*θ*_ = (*α, β, ν, μ*_0_) will be the parameters of a Normal-Inverse Gamma distribution.

Notice that the state sequence does not depend on any prior. Though there are several schemes available to sample **q**, [58] has argued strongly in favor of Forward-Backward sampling [57], which yields Forward-Backward Gibbs sampling (FBG) above. Variations of this have been implemented for segmentation of aCGH data before [60,62,78]. However, since in each iteration a quadratic number of terms has to be calculated at each position to obtain the forward variables, and a state has to be sampled at each position in the backward step, this method is still expensive for large data. Recently, [62] have introduced *compressed FBG* by sampling over a shorter sequence of sufficient statistics of data segments which are likely to come from the same underlying state. Let 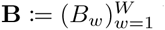 be a partition of **y** into *W* blocks. Each block *B_w_* contains *n_w_* elements. Let *y_w_*,_*k*_ the *k*-th element in *B_w_*. The forward variable *α_w_*(*j*) for this block needs to take into account the *n_w_* emissions, the transitions into state *j*, and the *n_w_* – 1 self-transitions, which yields

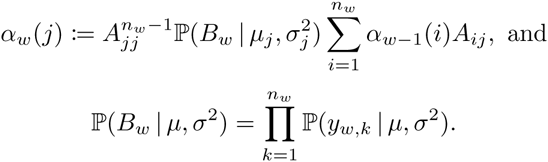

The ideal block structure would correspond to the actual, unknown segmentation of the data. Any subdivision thereof would decrease the compression ratio, and thus the speedup, but still allow for recovery of the true breakpoints. In addition, such a segmentation would yield sufficient statistics for the likelihood computation that corresponds to the true parameters of the state generating a segment. Using wavelet theory, we show that such a block structure can be easily obtained.

### Wavelet theory preliminaries

Here, we review some essential wavelet theory; for details, see [79,80]. Let

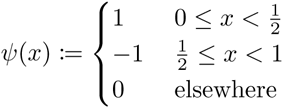

be the Haar wavelet [81], and *ψ*_*j*,*k*_(*x*) := 2^*j*/2^*ψ*(2^*j*^*x* – *k*); *j* and *k* are called the *scale* and *shift* parameter. Any square-integrable function over the unit interval, *f* ∈ *L*^2^([0,1)), can be approximated using the orthonormal basis {*ψ*_*j*,*k*_ | *j, k* ∈ ℤ, –1 ≤ *j*, 0 ≤ *k* ≤ 2^*j*^ – 1 }, admitting a *multiresolution analysis* [82,83]. Essentially, this allows us to express a function *f*(*x*) using scaled and shifted copies of one simple basis function *ψ*(*x*) which is spatially localized, i. e. non-zero on only a finite interval in *x*. The Haar basis is particularly suited for expressing piecewise constant functions.

Finite data **y** := (*y*_0_,…, *y*_*T*–1_) can be treated as an equidistant sample *f*(*x*) by scaling the indices to the unit interval using 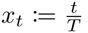. Let *h* := log_2_*T*. Then **y** can be expressed exactly as a linear combination over the Haar wavelet basis above, restricted to the maximum level of sampling resolution (*j* ≤ *h* – 1),

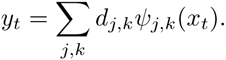

The *wavelet transform* **d** = ***W*****y** is an orthogonal endomorphism, and thus incurs neither redundancy nor loss of information. Surprisingly, **d** can be computed in linear time using the *pyramid algorithm* [82].

### Compression via wavelet shrinkage

The Haar wavelet transform has an important property connecting it to block creation: Let **d̂** be a vector obtained by setting elements of **d** to zero, then 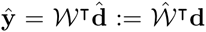 is called *selective wavelet reconstruction* (SW). If all coefficients *d*_*j,k*_ with *ψ*_*j,k*_(*x_t_*) ≠ *ψ*_*j,k*_(*x*_*t*+1_) are set to zero for some *t*, then *ŷ_t_* = *ŷ*_*t*+1_, which implies a block structure on **ŷ**. Conversely, blocks of size > 2 (to account for some pathological cases) can only be created using SW. *This general equivalence between SW and compression is central to our method*. Note that **ŷ** does *not* have to be computed explicitly; the block boundaries can be inferred from the position of zero-entries in **d̂** alone.

Assume all HMM states had the same emission variance *σ*^2^. Since each state is associated with an emission mean, finding **q** can be viewed as a regression or smoothing problem of finding an estimate ***μ̂*** of a piecewise constant function *μ* whose range is precisely the set of emission means, i. e.

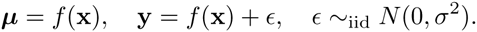

Unfortunately, regression methods typically do not limit the number of distinct values recovered, and will instead return some estimate **ŷ** ≠ ***μ̂***. However, if ***ŷ*** is piecewise constant and minimizes ∥***μ*** – **ŷ**∥, the sample means of each block are close to the true emission means. This yields high likelihood for their corresponding state and hence a strong posterior distribution, leading to fast convergence. Furthermore, the change points in **μ** must be close to change points in **ŷ**, since moving block boundaries incurs additional loss, allowing for approximate recovery of true breakpoints. **ŷ** might however induce additional block boundaries that reduce the compression ratio.

In a series of ground-breaking papers, Donoho, Johnstone *et al*. [84–88] showed that SW could in theory be used as an almost ideal spatially adaptive regression method. Assuming one could provide an oracle Δ(**μ**, **y**) that would know the true ***μ***, then there exists a method 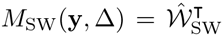 using an optimal subset of wavelet coefficients provided by **Δ** such that the quadratic risk of 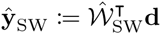 is bounded as

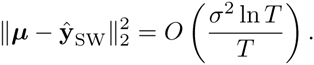

By definition, *M*_sw_ would be the best compression method under the constraints of the Haar wavelet basis. This bound is generally unattainable, since the oracle cannot be queried. Instead, they have shown that for a method *M*_WCT_(**y**, λ*σ*) called *wavelet coefficient thresholding*, which sets coefficients to zero whose absolute value is smaller than some threshold λ*σ*, there exists some 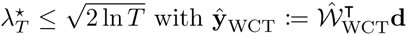 such that

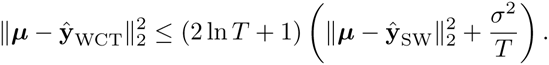

This 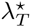 is minimax, i. e. the maximum risk incured over all possible data sets is not larger than that of any other threshold, and no better bound can be obtained. It is not easily computed, but for large *T*, on the order of tens to hundreds, the *universal threshold* 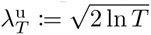 is asymptotically minimax. In other words, for data large enough to warrant compression, universal thresholding is the best method to approximate ***μ***, and thus the best wavelet-based compression scheme for a given noise level *σ*^2^.

### Integrating wavelet shrinkage into FBG

This compression method can easily be extended to multiple emission variances. Since we use a thresholding method, decreasing the variance simply subdivides existing blocks. If the threshold is set to the smallest emission variance among all states, **ŷ** will approximately preserve the breakpoints around those low-variance segments. Those of high variance are split into blocks of lower sample variance; see [89, 90] for an analytic expression. While the variances for the different states are not known, FBG provides *a priori* samples in each iteration. We hence propose the following simple adaptation: *In each sampling iteration, use the smallest sampled variance parameter to create a new block sequence via wavelet thresholding (Algorithm 1)*.

#### Algorithm 1 Dynamically adaptive FBG for HMMs

1. **procedure** HAMMLET(**y**, τ_***A***_, τ_*θ*_, τ_**π**_)
2. *T* ← |**y** |
3. 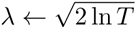
4. ***A*** ~ ℙ(***A*** |τ_***A***_)
5. *θ* ~ ℙ(*θ* | τ_*θ*_)
6. **π** ~ ℙ(**π** | τ_**π**_)
7. **for** *i* = 1,…,*N* **do**
8. 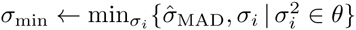
9. Create block sequence **B** from threshold λσ_min_
10. **q** ~ ℙ(**q** | ***A***, **B**, *θ, π*) using Forward-Backward sampling
11. Add count of marginal states for **q** to result
12. 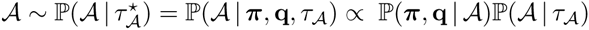
13. 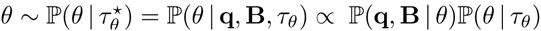
14. 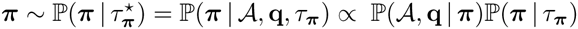
15. **end for**
16. **end procedure**

While intuitively easy to understand, provable guarantees for the optimality of this method, specifically the correspondence between the wavelet and the HMM domain remain an open research topic. A potential problem could arise if all sampled variances are too large. In this case, blocks would be under-segmented, yield wrong posterior variances and hide possible state transitions. As a safeguard against over-compression, we use the standard method to estimate the variance of constant noise in shrinkage applications,

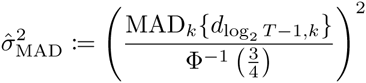

as an estimate of the variance in the dominant component, and modify the threshold definition to λ · min {σ̂_MAD_,*σ_i_* ∈ *θ* }. If the data is not i.i.d., 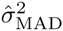 will systematically underestimate the true variance [28]. In this case, the blocks get smaller than necessary, thus decreasing the compression.

### A data structure for dynamic compression

The necessity to recreate a new block sequence in each iteration based on the most recent estimate of the smallest variance parameter creates the challenge of doing so with little computational overhead, specifically without repeatedly computing the inverse wavelet transform or considering all *T* elements in other ways. We achieve this by creating a simple tree-based data structure.

The pyramid algorithm yields **d** sorted according to (*j, k*). Again, let *h* := log_2_*T*, and *ℓ* := *h* – *j*. We can map the wavelet *ψ*_*j*,*k*_ to a perfect binary tree of height *h* such that all wavelets for scale *j* are nodes on level *i*, nodes within each level are sorted according to *k*, and *i* is increasing from the leaves to the root (Fig. 7). **d** represents a breadth-first search (BFS) traversal of that tree, with *d*_*j*,*k*_ being the entry at position [2^*j*^ + *k*. Adding *y_i_* as the *i*-th leaf on level *ℓ* = 0, each non-leaf node represents a wavelet which is non-zero for the *n* := 2^*ℓ*^ data points *y_t_*, for *t* in the the interval *I*_*j,k*_ := [*kn*, (*k* + 1)*n* – 1] stored in the leaves below; notice that for the leaves, *kn* = *t*.

**Figure 7.**
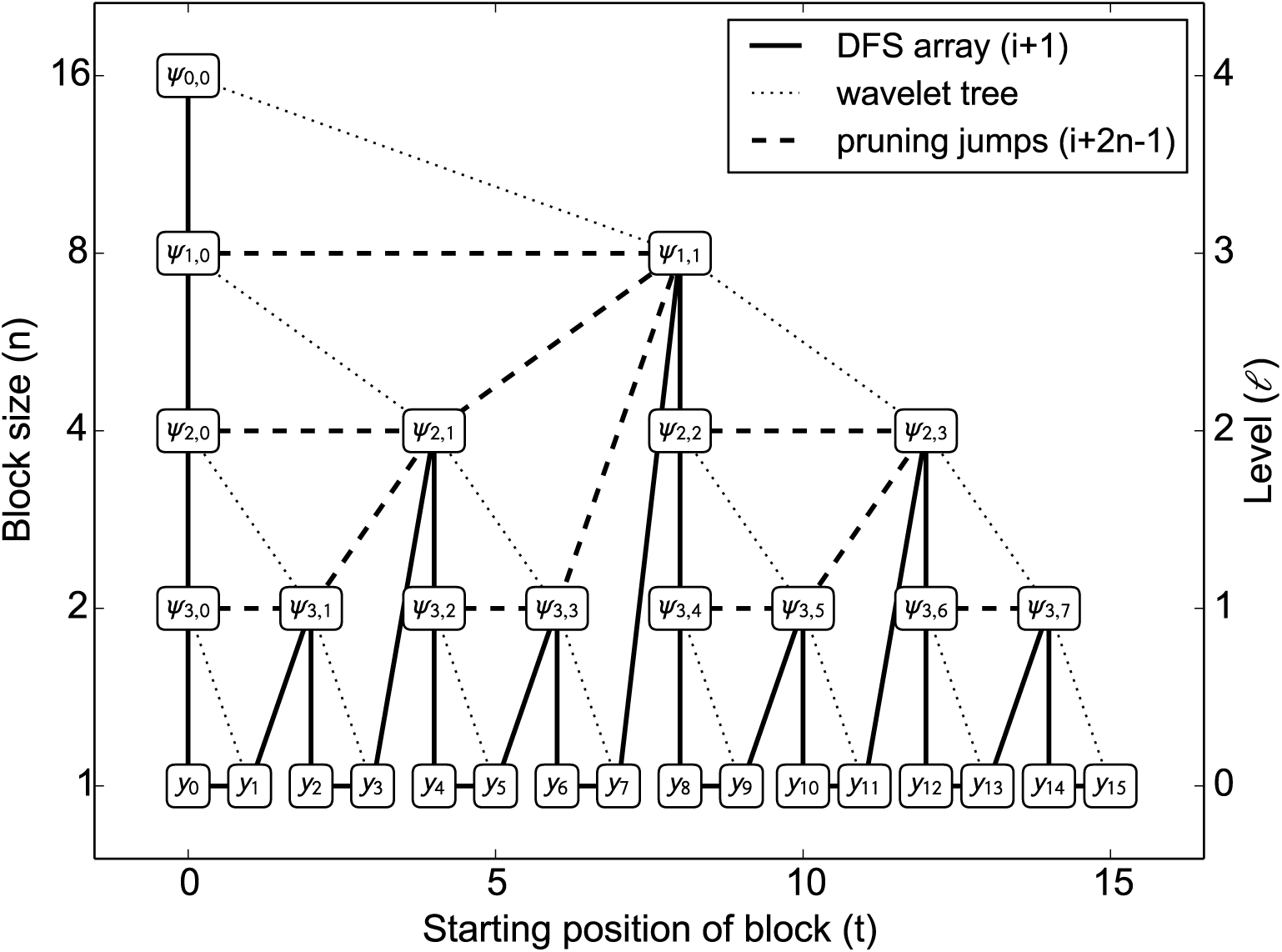
Mapping of wavelets *ψ*_*j*,*k*_ and data points *y_t_* to tree nodes *N*_*ℓ,t*_. Each node is the root of a subtree with *n* = 2^*ℓ*^ leaves; pruning that subtree yields a block of size *n*, starting at position *t*. For instance, the node *N*_1,e_ is located at position 13 of the DFS array (solid line), and corresponds to the wavelet *ψ*_3,3_. A block of size *n* = 2 can be created by pruning the subtree, which amounts to advancing by 2*n* – 1 = 3 positions (dashed line), yielding *N*_3,8_ at position 16, which is the wavelet *ψ*_1,1_. Thus the number of steps for creating blocks per iteration is at most the number of nodes in the tree, and thus strictly smaller than 2*T*.

This implies that the leaves in any subtree all have the same value after wavelet thresholding if all the wavelets in this subtree are set to zero. We can hence avoid computing the inverse wavelet transform to create blocks. Instead, each node stores the maximum absolute wavelet coefficient in the entire subtree, as well as the sufficient statistics required for calculating the likelihood function. More formally, a node *N*_*ℓ*,*t*_ corresponds to wavelet *ψ*_*j*,*k*_, with *ℓ* = *h* – *j* and *t* = *k*2^*ℓ*^ (*ψ*_–1,0_ is simply constant on the [0,1) interval and has no effect on block creation, thus we discard it). Essentially, *ℓ* numbers the levels beginning at the leaves, and *t* marks the start position of the block when pruning the subtree rooted at *N*_*ℓ*,*t*_. The members stored in each node are:

- The number of leaves, corresponding to the block size:

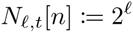
- The sum of data points stored in the subtree leaves:

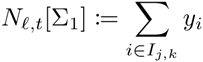
- Similarly, the sum of squares:

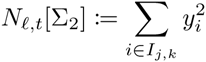
- The maximum absolute wavelet coefficient of the subtree, including the current *d*_*j*,*k*_ itself:

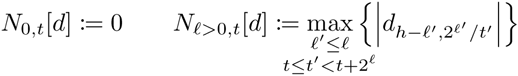

All these values can be computed recursively from the child nodes in linear time. As some real data sets contain salt-and-pepper noise, which manifests as isolated large coefficients on the lowest level, its is possible to ignore the first level in the maximum computation so that no information to create a single-element block for outliers is passed up the tree. We refer to this technique as *noise control*. Notice that this does not imply that blocks are only created at even *t*, since true transitions manifest in coefficients on multiple levels.

The block creation algorithm is simple: upon construction, the tree is converted to *depth-first search* (DFS) order, which simply amounts to sorting the BFS array according to (*kn, j*), and can be performed using linear-time algorithms such as radix sort; internally, we implemented a different linear-time implementation mimicking tree traversal using a stack. Given a threshold, the tree is then traversed in DFS order by iterating linearly over the array (Fig. 7, solid lines). Once the maximum coefficient stored in a node is less or equal to the threshold, a block of size *n* is created, and the entire subtree is skipped (dashed lines). As the tree is perfect binary and complete, the next array position in DFS traversal after pruning the subtree rooted at the node at index *i* is simply obtained as *i* + 2*n* – 1, so no expensive pointer structure needs to be maintained, leaving the tree data structure a simple flat array. An example of dynamic block creation is given in Fig. 8.

**Figure 8.**
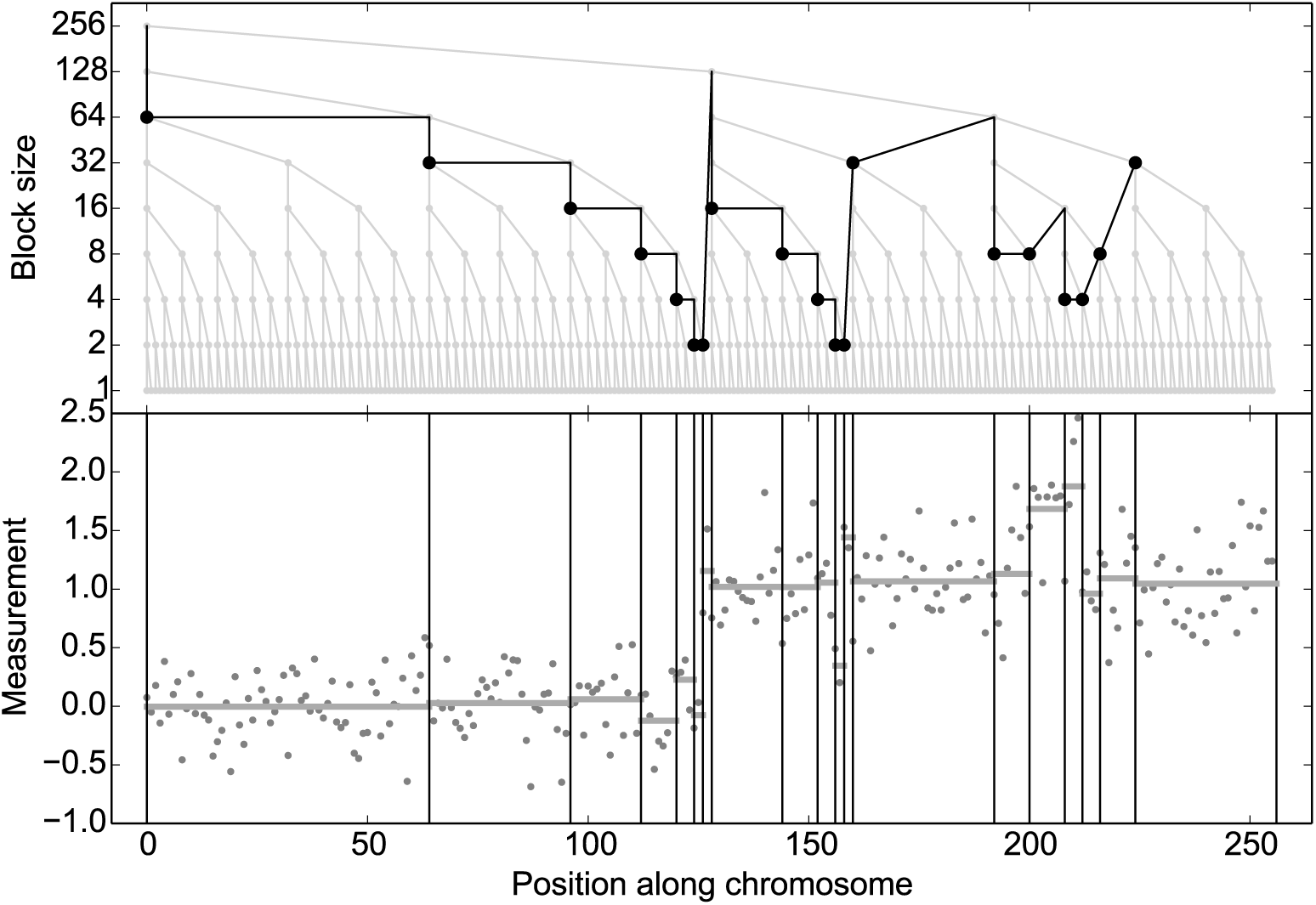
Example of dynamic block creation. The data is of size *T* = 256, so the wavelet tree contains 512 nodes. Here, only 37 entries had to be checked against the threshold (dark line), 19 of which (round markers) yielded a block (vertical lines on the bottom). Sampling is hence done on a short array of 19 blocks instead of 256 individual values, thus the compression ratio is 13.5. The horizontal lines in the bottom subplot are the block means derived from the sufficient statistics in the nodes. Notice how the algorithm creates small blocks around the breakpoints, e. g. at *t* ≈ 125, which requires traversing to lower levels and thus induces some additional blocks in other parts of the tree (left subtree), since all block sizes are powers of 2. This somewhat reduces the compression ratio, which is unproblematic as it increases the degrees of freedom in the sampler.

Once the Gibbs sampler converges to a set of variances, the block structure is less likely to change. To avoid recreating the same block structure over and over again, we employ a technique called *block structure prediction*. Since the different block structures are subdivisions of each other that occur in a specific order for decreasing σ^2^, there is a simple bijection between the number of blocks and the block structure itself. Thus, for each block sequence length we register the minimum and maximum variance that creates that sequence. Upon entering a new iteration, we check if the current variance would create the same number of blocks as in the previous iteration, which guarantees that we would obtain the same block sequence, and hence can avoid recomputation.

The wavelet tree data structure can be readily extended to multivariate data of dimensionality *m*. Instead of storing *m* different trees and reconciling *m* different block patterns in each iteration, one simply stores *m* different values for each sufficient statistic in a tree node. Since we have to traverse into the combined tree if the coefficient of any of the m trees was below the threshold, we simply store the largest *N*_*ℓ, t*_[*d*] among the corresponding nodes of the trees, which means that the block creation can be done in *O*(*T*) instead of *O*(*mT*), i. e. the dimensionality of the data only enters into the creation of the data structure, but not the query during sampling iterations.

### Automatic priors

While Bayesian methods allow for inductive bias such as the expected location of means, it is desirable to be able to use our method even when little domain knowledge exists, or large variation is expected, such as the lab and batch effects commonly observed in micro-arrays [91], as well as unknown means due to sample contamination. Since FBG does require a prior even in that case, we propose the following method to specify hyperparameters of a weak prior automatically. Posterior samples of means and variances are drawn from a Normal-Inverse Gamma distribution (*μ, σ*^2^) ~ NIΓ(*μ*_0_, *ν, α, β*), whose marginals simply separate into a Normal and an Inverse Gamma distribution

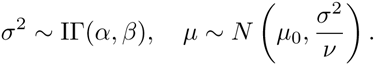

Let s^2^ be a user-defined variance (or automatically infered, e.g. from the largest of the finest detail coefficients, or 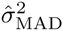), and *p* the desired probability to sample a variance not larger than *s*^2^. From the CDF of Ir we obtain

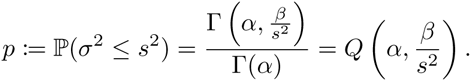

Ir has a mean for *α* > 1, and closed-form solutions for *α* ∈ ℕ. Furthermore, IΓ has positive skewness for *α* > 3. We thus let *α* = 2, which yields

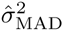

where *W*_–1_ is the negative branch of the Lambert *W*-function, which is transcendental. However, an excellent analytical approximation with a maximum error of 0.025% is given in [92], which yields

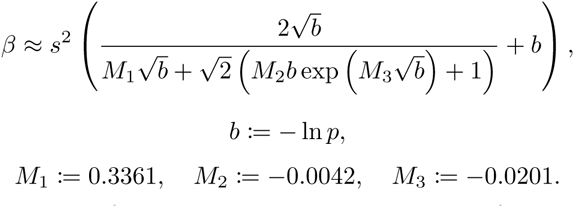

Since the mean of IΓ is 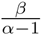, the expected variance of *μ* is 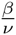 for *α* = 2. To ensure proper mixing, we could simply set 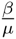 to the sample variance of the data, which can be estimated from the sufficient statistics in the root of the wavelet tree (the first entry in the array), provided that ***μ*** contained all states in almost equal number. However, due to possible class imbalance, means for short segments far away from *μ*_0_ can have low sampling probability, as they do not contribute much to the sample variance of the data. We thus define *δ* to be the sample variance of block means in the compression obtained by 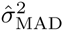, and take the maximum of those two variances. We thus obtain

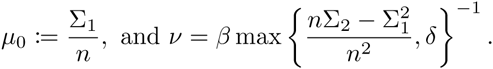

### Numerical issues

To assure numerical stability when working with probabilities, many HMM implementations resort to log-space computations, which involves a considerable number of expensive function calls (exp, log, pow); for instance, on Intel’s Nehalem architecture, log (FYL2X) requires 55 operations as opposed to 1 for adding and multiplying floating point numbers (FADD, FMUL) [93]. Our implementation, which differs from [62] greatly reduces the number of such calls by utilizing the block structure: The term accounting for emissions and self-transitions within the block can be written as

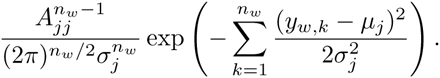

Any constant cancels out during normalization. Furthermore, exponentiation of potentially small numbers causes underflows. We hence move those terms into the exponent, utilizing the much stabler logarithm function.

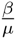

Using the block’s sufficient statistics

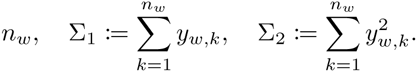

the exponent can be rewritten as

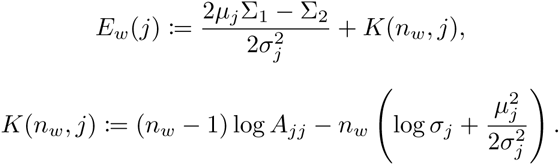

*K*(*n_w_*, *j*) can be precomputed for each iteration, thus greatly reducing the number of expensive function calls. Notice that the expressions above correspond to the canonical exponential family form exp(〈*t*(*x*), *θ*〉–*F*(*θ*)+*k*(*x*)) of a product of Gaussian distributions. Hence, equivalent terms can easily be derived for non-Gaussian emissions, implying that the same optimizations can be used in the general case of exponential family distributions: Only the dot product of the sufficient statistics *t*(*x*) and the parameters *θ* has to be computed in each iteration and for each block, while the log-normalizer *F*(*θ*) can be precomputed for each iteration, and the carrier measure *k*(*x*) (which is 0 for Gaussian emissions) only has to be computed once.

To avoid overflow of the exponential function, we subtract the largest such exponents among all states, hence *E_w_*(*j*) ≤ 0. This is equivalent to dividing the forward variables by

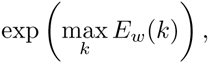

which cancels out during normalization. Hence we obtain

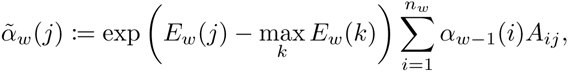

which are then normalized to

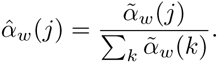

## Acknowledgements

We would like to thank Md Pavel Mahmud for helpful advice.

## Supporting information

The supplemental figures, our simulation data and results are available for download at https://zenodo.org/record/46263(DOI: 10.5281/zenodo.46263) [94], and are referenced as S1–S7 throughout the text. Additionally, the figures can also be viewed through our website at http://schlieplab.org/Supplements/HaMMLET/ for convenience. The implementation of HaMMLET and scripts to reproduce the simulation and evaluation are available at https://github.com/wiedenhoeft/HaMMLET/tree/biorxiv, and a snapshot is archived at https://zenodo.org/record/46262 (DOI: 10.5281/zenodo.46262) [95]. The high-density aCGH data [68] is available from http://www.ncbi.nlm.nih.gov/geo/query/acc.cgi?acc=GSE23949 (accession GEO:GSE23949). Coriell etc. data [77] is available from http://www.ncbi.nlm.nih.gov/geo/query/acc.cgi?acc=GSE16 (accession GEO:GSE16). The simulations of [66] are available from the original authors’ website at http://www.cbs.dtu.dk/~hanni/aCGH/. Notice that due to the use of a random number generator by HaMMLET, CBS and our simulations, individual results will differ slightly from the data provided in the supplement.

## References

1. Iafrate AJ, Feuk L, Rivera MN, Listewnik ML, Donahoe PK, Qi Y, et al. Detection of large-scale variation in the human genome. Nature Genetics. 2004 Sep;36(9):949–51. Available from: http://www.nature.com/ng/journal/v36/n9/full/ng1416.html.

2. Feuk L, Marshall CR, Wintle RF, Scherer SW. Structural variants: changing the landscape of chromosomes and design of disease studies. Human Molecular Genetics. 2006 Apr;15 Spec No:R57–66. Available from: http://www.ncbi.nlm.nih.gov/pubmed/16651370.

3. McCarroll SA, Altshuler DM. Copy-number variation and association studies of human disease. Nature Genetics. 2007 Jul;39(7 Suppl):S37–42. Available from: http://www.ncbi.nlm.nih.gov/pubmed/17597780.

4. Wain LV, Armour JAL, Tobin MD. Genomic copy number variation, human health, and disease. Lancet. 2009 Jul;374(9686):340–350. Available from: http://www.sciencedirect.com/science/article/pii/S014067360960249X.

5. Beckmann JS, Sharp AJ, Antonarakis SE. CNVs and genetic medicine (excitement and consequences of a rediscovery). Cytogenetic and Genome Research. 2008 Jan;123(1-4):7–16. Available from: http://www.karger.com/Article/FullText/184687.

6. Cook EH, Scherer SW. Copy-number variations associated with neuropsychiatric conditions. Nature. 2008 Oct;455(7215):919–23. Available from: http://dx.doi.org/10.1038/nature07458.

7. Buretic-Tomljanovic A, Tomljanovic D. Human genome variation in health and in neuropsychiatric disorders. Psychiatria Danubina. 2009 Dec;21(4):562–9. Available from: http://www.ncbi.nlm.nih.gov/pubmed/19935494.

8. Merikangas AK, Corvin AP, Gallagher L. Copy-number variants in neurodevelopmental disorders: promises and challenges. Trends in Genetics. 2009 Dec;25(12):536–44. Available from: http://www.sciencedirect.com/science/article/pii/S016895250900211X.

9. Cho EK, Tchinda J, Freeman JL, Chung YJ, Cai WW, Lee C. Array-based comparative genomic hybridization and copy number variation in cancer research. Cytogenetic and Genome Research. 2006 Jan;115(3-4):262–72. Available from: http://www.karger.com/Article/FullText/95923.

10. Shlien A, Malkin D. Copy number variations and cancer susceptibility. Current Opinion in Oncology. 2010 Jan;22(1):55–63. Available from: http://www.ncbi.nlm.nih.gov/pubmed/19952747.

11. Feuk L, Carson AR, Scherer SW. Structural variation in the human genome. Nature Reviews Genetics. 2006 Feb;7(2):85–97. Available from: http://www.nature.com/nrg/journal/v7/n2/full/nrg1767.html.

12. Beckmann JS, Estivill X, Antonarakis SE. Copy number variants and genetic traits: closer to the resolution of phenotypic to genotypic variability. Nature Reviews Genetics. 2007 Aug;8(8):639–46. Available from: http://www.ncbi.nlm.nih.gov/pubmed/17637735.

13. Sharp AJ. Emerging themes and new challenges in defining the role of structural variation in human disease. Human Mutation. 2009 Feb;30(2):135–44. Available from: http://www.ncbi.nlm.nih.gov/pubmed/18837009.

14. Olshen AB, Venkatraman ES, Lucito R, Wigler M. Circular binary segmentation for the analysis of array-based DNA copy number data. Biostatistics (Oxford, England). 2004 Oct;5(4):557–572. Available from: http://biostatistics.oxfordjournals.org/content/5/4/557.short.

15. Venkatraman ES, Olshen AB. A faster circular binary segmentation algorithm for the analysis of array CGH data. Bioinformatics. 2007 Mar;23(6):657–63. Available from: http://bioinformatics.oxfordjournals.org/content/23/6/657.short.

16. Xing B, Greenwood CMT, Bull SB. A hierarchical clustering method for estimating copy number variation. Biostatistics (Oxford, England). 2007 Jul;8(3):632–53. Available from: http://biostatistics.oxfordjournals.org/content/8/3/632.full.

17. Fridlyand J, Snijders AM, Pinkel D, Albertson DG, Jain AN. Hidden Markov models approach to the analysis of array CGH data. Journal of Multivariate Analysis. 2004 Jul;90(1):132–153. Available from: http://www.sciencedirect.com/science/article/pii/S0047259X04000260.

18. Garnis C, Coe BP, Zhang L, Rosin MP, Lam WL. Overexpression of LRP12, a gene contained within an 8q22 amplicon identified by high-resolution array CGH analysis of oral squamous cell carcinomas. Oncogene. 2004 Apr;23(14):2582–6. Available from: http://www.ncbi.nlm.nih.gov/pubmed/14676824.

19. Albertson DG, Ylstra B, Segraves R, Collins C, Dairkee SH, Kowbel D, et al. Quantitative mapping of amplicon structure by array CGH identifies CYP24 as a candidate oncogene. Nature Genetics. 2000 Jun;25(2):144–6. Available from: http://www.ncbi.nlm.nih.gov/pubmed/10835626.

20. Veltman JA, Fridlyand J, Pejavar S, Olshen AB, Korkola JE, DeVries S, et al. Array-based comparative genomic hybridization for genome-wide screening of DNA copy number in bladder tumors. Cancer Research. 2003 Jun;63(11):2872–80. Available from: http://www.ncbi.nlm.nih.gov/pubmed/12782593.

21. Autio R, Hautaniemi S, Kauraniemi P, Yli-Harja O, Astola J, Wolf M, et al. CGH-Plotter: MATLAB toolbox for CGH-data analysis. Bioinformatics. 2003 Sep;19(13):1714–1715. Available from: http://bioinformatics.oxfordjournals.org/content/19/13/1714.

22. Pollack JR, Sørlie T, Perou CM, Rees CA, Jeffrey SS, Lonning PE, et al. Microarray analysis reveals a major direct role of DNA copy number alteration in the transcriptional program of human breast tumors. Proceedings of the National Academy of Sciences of the United States of America. 2002 Oct;99(20):12963–8. Available from: http://www.pnas.org/content/99/20/12963.long.

23. Eilers PHC, de Menezes RX. Quantile smoothing of array CGH data. Bioinformatics. 2005 Apr;21(7):1146–53. Available from: http://bioinformatics.oxfordjournals.org/content/21/7/1146.full.

24. Cleveland WS. Robust Locally Weighted Regression and Smoothing Scatterplots. Journal of the American Statistical Association. 1979 Apr;74(368). Available from: http://www.tandfonline.com/doi/abs/10.1080/01621459.1979.10481038.

25. Beheshti B, Braude I, Marrano P, Thorner P, Zielenska M, Squire JA. Chromosomal localization of DNA amplifications in neuroblastoma tumors using cDNA microarray comparative genomic hybridization. Neoplasia. 2003 Jan;5(1):53–62. Available from: http://www.ncbi.nlm.nih.gov/pmc/articles/PMC1502121/.

26. Polzehl J, Spokoiny VG. Adaptive weights smoothing with applications to image restoration. Journal of the Royal Statistical Society: Series B (Statistical Methodology). 2000 May;62(2):335–354. Available from: http://doi.wiley.com/10.1111/1467-9868.00235.

27. Hupe P, Stransky N, Thiery JP, Radvanyi F, Barillot E. Analysis of array CGH data: from signal ratio to gain and loss of DNA regions. Bioinformatics. 2004 Dec;20(18):3413–22. Available from: http://bioinformatics.oxfordjournals.org/content/20/18/3413.

28. Wang Y, Wang S. A novel stationary wavelet denoising algorithm for array-based DNA Copy Number data. International Journal of Bioinformatics Research and Applications. 2007;3(x):206–222.

29. Hsu L, Self SG, Grove D, Randolph T, Wang K, Delrow JJ, et al. Denoising array-based comparative genomic hybridization data using wavelets. Biostatistics (Oxford, England). 2005 Apr;6(2):211–26. Available from: http://biostatistics.oxfordjournals.org/content/6/2/211.

30. Nguyen N, Huang H, Oraintara S, Vo A. A New Smoothing Model for Analyzing Array CGH Data. In: Proceedings of the 7th IEEE International Conference on Bioinformatics and Bioengineering. Boston, MA; 2007. Available from: http://ieeexplore.ieee.org/xpls/abs_all.jsp?arnumber=4375683.

31. Nguyen N, Huang H, Oraintara S, Vo A. Stationary wavelet packet transform and dependent Laplacian bivariate shrinkage estimator for array-CGH data smoothing. Journal of Computational Biology. 2010 Feb;17(2):139–52. Available from: http://online.liebertpub.com/doi/abs/10.1089/cmb.2009.0013.

32. Huang H, Nguyen N, Oraintara S, Vo A. Array CGH data modeling and smoothing in Stationary Wavelet Packet Transform domain. BMC Genomics. 2008 Jan;9 Suppl 2:S17. Available from: http://www.ncbi.nlm.nih.gov/pmc/articles/PMC2559881/.

33. Holt C, Losic B, Pai D, Zhao Z, Trinh Q, Syam S, et al. WaveCNV: allele-specific copy number alterations in primary tumors and xenograft models from next-generation sequencing. Bioinformatics. 2013 Nov;p. btt611-. Available from:http://bioinformatics.oxfordjournals.org/content/early/2013/11/26/bioinformatics.btt611.full.

34. Price TS, Regan R, Mott R, Hedman A, Honey B, Daniels RJ, et al. SW-ARRAY: a dynamic programming solution for the identification of copy-number changes in genomic DNA using array comparative genome hybridization data. Nucleic Acids Research. 2005 Jan;33(11):3455–3464. Available from: http://nar.oxfordjournals.org/content/33/11/3455.

35. Tsourakakis CE, Peng R, Tsiarli MA, Miller GL, Schwartz R. Approximation algorithms for speeding up dynamic programming and denoising aCGH data. Journal of Experimental Algorithmics. 2011 May;16:1.1. Available from: http://dl.acm.org/citation.cfm?id=1963190.2063517.

36. Olshen AB, Venkatraman ES. Change-point analysis of array-based comparative genomic hybridization data. ASA Proceedings of the Joint Statistical Meetings. 2002;p. 2530–2535.

37. Picard F, Robin S, Lavielle M, Vaisse C, Daudin JJ. A statistical approach for array CGH data analysis. BMC Bioinformatics. 2005 Jan;6(1):27. Available from: http://www.biomedcentral.com/1471-2105/6/27.

38. Myers CL, Dunham MJ, Kung SY, Troyanskaya OG. Accurate detection of aneuploidies in array CGH and gene expression microarray data. Bioinformatics. 2004 Dec;20(18):3533–43. Available from: http://bioinformatics.oxfordjournals.org/content/20/18/3533.

39. Wang P, Kim Y, Pollack J, Narasimhan B, Tibshirani R. A method for calling gains and losses in array CGH data. Biostatistics (Oxford, England). 2005 Jan;6(1):45–58. Available from: http://biostatistics.oxfordjournals.org/content/6/1/45.

40. Chen CH, Lee HCC, Ling Q, Chen HR, Ko YA, Tsou TS, et al. An allstatistics, high-speed algorithm for the analysis of copy number variation in genomes. Nucleic Acids Research. 2011 Jul;39(13):e89. Available from: http://nar.oxfordjournals.org/content/39/13/e89.full.

41. Jong K, Marchiori E, van der Vaart A, Ylstra B, Weiss M, Meijer G. Chromosomal Breakpoint Detection in Human Cancer. vol. 2611 of Lecture Notes in Computer Science. Cagnoni S, Johnson CG, Cardalda JJR, Marchiori E, Corne DW, Meyer JA, et al., editors. Berlin, Heidelberg: Springer; 2003. Available from: http://link.springer.com/10.1007/3-540-36605-9.

42. Baum LE, Petrie T. Statistical Inference for Probabilistic Functions of Finite State Markov Chains. The Annals of Mathematical Statistics. 1966;37(6):1554–1563. Available from: http://www.jstor.org/stable/2238772.

43. Snijders AM, Fridlyand J, Mans DA, Segraves R, Jain AN, Pinkel D, et al. Shaping of tumor and drug-resistant genomes by instability and selection. Oncogene. 2003 Jul;22(28):4370–9. Available from: http://www.ncbi.nlm.nih.gov/pubmed/12853973.

44. Sebat J, Lakshmi B, Troge J, Alexander J, Young J, Lundin P, et al. Large-scale copy number polymorphism in the human genome. Science. 2004 Jul;305(5683):525–8. Available from: http://www.sciencemag.org/content/305/5683/525.full.

45. Sebat J, Lakshmi B, Malhotra D, Troge J, Lese-Martin C, Walsh T, et al. Strong association of de novo copy number mutations with autism. Science. 2007 Apr;316(5823):445–449. Available from: http://www.sciencemag.org/content/316/5823/445.abstract.

46. Zhao X. An Integrated View of Copy Number and Allelic Alterations in the Cancer Genome Using Single Nucleotide Polymorphism Arrays. Cancer Research. 2004 May;64(9):3060–3071. Available from: http://cancerres.aacrjournals.org/content/64/9/3060.long.

47. de Vries BBA, Pfundt R, Leisink M, Koolen DA, Vissers LELM, Janssen IM, et al. Diagnostic genome profiling in mental retardation. American Journal of Human Genetics. 2005 Oct;77(4):606–616. Available from: http://www.sciencedirect.com/science/article/pii/S0002929707610088.

48. Nannya Y, Sanada M, Nakazaki K, Hosoya N, Wang L, Hangaishi A, et al. A robust algorithm for copy number detection using high-density oligonucleotide single nucleotide polymorphism genotyping arrays. Cancer Research. 2005 Jul;65(14):6071–9. Available from:http://cancerres.aacrjournals.org/content/65/14/6071.full.

49. Marioni JC, Thorne NP, Tavare S. BioHMM: a heterogeneous hidden Markov model for segmenting array CGH data. Bioinformatics. 2006 May;22(9):1144–6. Available from: http://bioinformatics.oxfordjournals.org/content/22/9/1144.long.

50. Korbel JO, Urban AE, Grubert F, Du J, Royce TE, Starr P, et al. Systematic prediction and validation of breakpoints associated with copy-number variants in the human genome. Proceedings of the National Academy of Sciences of the United States of America. 2007 Jun;104(24):10110–10115. Available from: http://www.pnas.org/content/104/24/10110.full.

51. Cahan P, Godfrey LE, Eis PS, Richmond TA, Selzer RR, Brent M, et al. wuHMM: a robust algorithm to detect DNA copy number variation using long oligonucleotide microarray data. Nucleic Acids Research. 2008 Apr;36(7):e41. Available from: http://nar.oxfordjournals.org/content/36/7/e41.full.

52. Rueda OM, Diaz-Uriarte R. RJaCGH: Bayesian analysis of aCGH arrays for detecting copy number changes and recurrent regions. Bioinformatics. 2009 Aug;25(15):1959–1960. Available from: http://www.ncbi.nlm.nih.gov/pmc/articles/PMC2712338/.

53. Bilmes J. A Gentle Tutorial of the EM Algorithm and its Application to Parameter Estimation for Gaussian Mixture and Hidden Markov Models; 1998. Available from: http://citeseerx.ist.psu.edu/viewdoc/summary?doi=10.1.1.28.613.

54. Rabiner LR. A tutorial on Hidden Markov Models and Selected Applications in Speech Recognition. Proceedings of the IEEE. 1989;77:257–286. Available from: http://ieeexplore.ieee.org/lpdocs/epic03/wrapper.htm?arnumber=18626.

55. Viterbi A. Error bounds for convolutional codes and an asymptotically optimum decoding algorithm. IEEE Transactions on Information Theory. 1967 Apr;13(2):260–269. Available from: http://ieeexplore.ieee.org/lpdocs/epic03/wrapper.htm?arnumber=1054010.

56. Forney GD. The Viterbi algorithm. Proceedings of the IEEE. 1973;61(3):268–278. Available from: http://ieeexplore.ieee.org/lpdocs/epic03/wrapper.htm?arnumber=1450960.

57. Chib S. Calculating posterior distributions and modal estimates in Markov mixture models. Journal of Econometrics. 1996;75(1):79–97. Available from: http://www.sciencedirect.com/science/article/pii/0304407695017704.

58. Scott SL. Bayesian Methods for Hidden Markov Models: Recursive Computing in the 21st Century. Journal of the American Statistical Association. 2002;97(457):337–351. Available from: http://www.jstor.org/stable/3085787.

59. Guha S, Li Y, Neuberg D. Bayesian Hidden Markov Modeling of Array CGH Data. Harvard University Biostatistics Working Paper Series. 2006;(24). Available from: http://biostats.bepress.com/harvardbiostat/paper24.

60. Shah SP, Xuan X, DeLeeuw RJ, Khojasteh M, Lam WL, Ng R, et al. Integrating copy number polymorphisms into array CGH analysis using a robust HMM. Bioinformatics. 2006 Jul;22(14):e431–9. Available from: http://bioinformatics.oxfordjournals.org/content/22/14/e431.

61. Shah SP, Lam WL, Ng RT, Murphy KP. Modeling recurrent DNA copy number alterations in array CGH data. Bioinformatics. 2007 Jul;23(13):i450–i458. Available from: http://bioinformatics.oxfordjournals.org/content/23/13/i450.short.

62. Mahmud MP, Schliep A. Fast MCMC sampling for Hidden Markov Models to determine copy number variations. BMC Bioinformatics. 2011 Jan;12:428. Available from: http://www.biomedcentral.com/1471-2105/12/428.

63. Ben-Yaacov E, Eldar YC. A fast and flexible method for the segmentation of aCGH data. Bioinformatics. 2008 Aug;24(16):i139–45. Available from: http://bioinformatics.oxfordjournals.org/content/24/16/i139.long.

64. Wang J, Meza-Zepeda LA, Kresse SH, Myklebost O. M-CGH: analysing microarray-based CGH experiments. BMC Bioinformatics. 2004 Jun;5(1):74. Available from: http://www.biomedcentral.com/1471-2105/5/74.

65. Lai WR, Johnson MD, Kucherlapati R, Park PJ. Comparative analysis of algorithms for identifying amplifications and deletions in array CGH data. Bioinformatics. 2005 Oct;21(19):3763–3770. Available from: http://bioinformatics.oxfordjournals.org/content/21/19/3763.short.

66. Willenbrock H, Fridlyand J. A comparison study: applying segmentation to array CGH data for downstream analyses. Bioinformatics. 2005 Nov;21(22):4084–91. Available from: http://bioinformatics.oxfordjournals.org/content/21/22/4084.long.

67. Hodgson G, Hager JH, Volik S, Hariono S, Wernick M, Moore D, et al. Genome scanning with array CGH delineates regional alterations in mouse islet carcinomas. Nature Genetics. 2001 Dec;29(4):459–64. Available from: http://www.nature.com/ng/journal/v29/n4/full/ng771.html.

68. Edgren H, Murumagi A, Kangaspeska S, Nicorici D, Hongisto V, Kleivi K, et al. Identification of fusion genes in breast cancer by paired-end RNA-sequencing. Genome Biology. 2011 Jan;12(1):R6. Available from: http://genomebiology.com/2011/12/1/R6.

69. Burdall S, Hanby A, Lansdown M, Speirs V. Breast cancer cell lines: friend or foe? Breast Cancer Research. 2003;5(2):89–95. Available from: http://breast-cancer-research.com/content/5/2/89.

70. Holliday DL, Speirs V. Choosing the right cell line for breast cancer research. Breast Cancer Research. 2011 Jan;13(4):215. Available from: http://www.ncbi.nlm.nih.gov/pmc/articles/PMC3236329/.

71. Van Wieringen WN, Van De Wiel MA, Ylstra B. Weighted clustering of called array CGH data. Biostatistics (Oxford, England). 2008 Jul;9(3):484–500. Available from: http://biostatistics.oxfordjournals.org/content/9/3/484.full.

72. Liu J, Mohammed J, Carter J, Ranka S, Kahveci T, Baudis M. Distance-based clustering of CGH data. Bioinformatics (Oxford, England). 2006 Aug;22(16):1971–8. Available from: http://bioinformatics.oxfordjournals.org/content/22/16/1971.short.

73. Gonzalez JR, Subirana I, Escaramis G, Peraza S, Caceres A, Estivill X, et al. Accounting for uncertainty when assessing association between copy number and disease: a latent class model. BMC Bioinformatics. 2009 Jan;10(1):172. Available from:http://bmcbioinformatics.biomedcentral.com/articles/10.1186/1471-2105-10-172.

74. van de Wiel MA, van Wieringen WN. CGHregions: dimension reduction for array CGH data with minimal information loss. Cancer informatics. 2007 Jan;3:55–63. Available from:http://www.pubmedcentral.nih.gov/articlerender.fcgi?artid=2675846.

75. Yin Xl, Li J. A general graphical framework for detecting copy number variations. In: 8th Annual International Conference on Computational Systems Bioinformatics. Life Sciences Society; 2009. Available from: http://www.csb2009a.org/pdf/060Li.pdf.

76. (Dzgür A, (Dzgür L, Güngor T. Text Categorization with Class-Based and Corpus-Based Keyword Selection. Proceedings of the 20th International Conference on Computer and Information Sciences. 2005;3733:606–615. Available from: http://link.springer.com/chapter/10.1007/11569596_63.

77. Snijders AM, Nowak N, Segraves R, Blackwood S, Brown N, Conroy J, et al. Assembly of microarrays for genome-wide measurement of DNA copy number. Nature Genetics. 2001 Nov;29(3):263–4. Available from: http://dx.doi.org/10.1038/ng754.

78. Mahmud MP, Schliep A. Speeding up Bayesian HMM by the four Russians method. In: Proceedings of the 11th International Conference on Algorithms in Bioinformatics 2011. p. 188–200. Available from: http://dl.acm.org/citation.cfm?id=2039945.2039962.

79. Daubechies I. Ten Lectures on Wavelets; 1992. Available from: http://epubs.siam.org/doi/book/10.1137/1.9781611970104.

80. Mallat SG. A wavelet tour of signal processing: The Sparse Way. Burlington, MA: Academic Press; 2009. Available from: http://dl.acm.org/citation.cfm?id=1525499.

81. Haar A. Zur Theorie der orthogonalen Funktionensysteme. Mathematische Annalen. 1910 Sep;69(3):331–371. Available from: http://link.springer.com/10.1007/BF01456326.

82. Mallat SG. A theory for multiresolution signal decomposition: the wavelet representation. IEEE Transactions on Pattern Analysis and Machine Intelligence. 1989 Jul;11(7):674–693. Available from: http://ieeexplore.ieee.org/lpdocs/epic03/wrapper.htm?arnumber=192463.

83. Mallat SG. Multiresolution approximations and wavelet orthonormal bases of L^2^(R). Transactions of the American Mathematical Society. 1989 Jan;315(1):69–69. Available from: http://www.ams.org/tran/1989-315-01/S0002-9947-1989-1008470-5/.

84. Donoho DL, Johnstone IM. Ideal spatial adaptation by wavelet shrinkage. Biometrika. 1994 Sep;81(3):425–455. Available from: http://biomet.oxfordjournals.org/content/81/3/425.

85. Donoho DL, Johnstone IM. Asymptotic minimaxity of wavelet estimators with sampled data. Statistica Sinica. 1999;9:1–32. Available from: http://citeseerx.ist.psu.edu/viewdoc/summary?doi=10.1.1.184.5690.

86. Donoho DL, Johnstone IM. Minimax estimation via wavelet shrinkage. The Annals of Statistics. 1998;26(3):879–921. Available from: http://www.jstor.org/stable/120061.

87. Donoho DL, Johnstone IM. Threshold selection for wavelet shrinkage of noisy data. In: Proceedings of 16th Annual International Conference of the IEEE Engineering in Medicine and Biology Society. Baltimore, MD: IEEE; 1994. p. 24a–25a. Available from: http://ieeexplore.ieee.org/lpdocs/epic03/wrapper.htm?arnumber=412133.

88. Donoho DL, Johnstone IM, Kerkyacharian G, Picard D. Wavelet Shrinkage: Asymptopia? Journal of the Royal Statistical Society: Series B (Statistical Methodology). 1995;57(2):301–369. Available from: http://www.jstor.org/stable/2345967.

89. Percival DB, Mofjeld HO. Analysis of Subtidal Coastal Sea Level Fluctuations Using Wavelets. Journal of the American Statistical Association. 1997;92(439):868. Available from: http://www.jstor.org/stable/2965551.

90. Serroukh A, Walden AT, Percival DB. Statistical Properties and Uses of the Wavelet Variance Estimator for the Scale Analysis of Time Series. Journal of the American Statistical Association. 2012 Feb;95. Available from: http://amstat.tandfonline.com/doi/abs/10.1080/01621459.2000.10473913.

91. Luo J, Schumacher M, Scherer A, Sanoudou D, Megherbi D, Davison T, et al. A comparison of batch effect removal methods for enhancement of prediction performance using MAQC-II microarray gene expression data. The Pharmacogenomics Journal. 2010 Aug;10(4):278–91. Available from: http://www.ncbi.nlm.nih.gov/pmc/articles/PMC2920074/.

92. Barry DA, Parlange JY, Li L, Prommer H, Cunningham CJ, Stagnitti F. Analytical approximations for real values of the Lambert W-function. Mathematics and Computers in Simulation. 2000 Aug;53(1-2):95–103. Available from: http://www.sciencedirect.com/science/article/pii/S0378475400001725.

93. Fog A. Instruction tables: Lists of instruction latencies, throughputs and micro-operation breakdowns for Intel, AMD and VIA CPUs; 2016. Available from: http://www.agner.org/optimize/instruction_tables.pdf.

94. Wiedenhoeft J, Brugel E, Schliep A. HaMMLET - Supplemental Material; 2016. Available from: http://dx.doi.org/10.5281/zenodo.46263.

95. Wiedenhoeft J, Brugel E. HaMMLET 0.0.0-alpha.1; 2016. Available from: http://dx.doi.org/10.5281/zenodo.46262.

